# Scaling laws for group-constrained, subject-specific task fMRI analyses

**DOI:** 10.64898/2026.07.22.740076

**Authors:** Ruimin Gao, Anna A. Ivanova

## Abstract

Functional MRI provides a powerful way to characterize the architecture of the human brain. Yet individual brains often vary in the exact anatomic locations of functionally specific regions; as a result of this variability, traditional random-effects group analyses (RFX) often require unrealistically large sample sizes to converge onto stable results. Here, we explore the scaling patterns for an alternative fMRI analysis framework that accounts for inter-individual topographic variability while still capturing spatial similarities—group-constrained, subject-specific (GcSS) analysis. Using sixteen contrasts from four Human Connectome Project tasks (spanning language, social cognition, motor, and working memory), we first show that the GcSS approach yields much larger effect sizes than traditional RFX analyses. We then investigate the impact of sample size (N=10–450) on the estimation of three key GcSS outputs: (1) group probabilistic maps for the contrast of interest, (2) group-level parcels that serve as anatomical constraints for subsequent functional region of interest (fROI) definitions in individual subjects, and (3) effect sizes of fROI responses to task conditions. For most contrasts, the probabilistic maps show good reliability at N=100 (intraclass correlation, ICC=.75) and excellent reliability at N=200 (ICC=.90). Most group parcels also achieve substantial agreement at N=100 (Dice coefficient, DC=.80). Critically, once robust group parcels are established, the subject-specific portion of the analysis can proceed with much smaller sample sizes: even samples of N=10 participants yield accurate effect size estimates within subject-specific fROIs, detecting 80% of practically meaningful effects (effect size ≥ 0.2); with N=20, this increases to 90%. The scaling patterns we observed held not only in the cortex, but also in the cerebellum. Our results challenge the view that fMRI research requires large samples: once the broad region of interest is established (with N=100 or higher), fROI-based analyses can yield generalizable results with small sample sizes (N=10 or N=20).

## Introduction

Functional specialization is a central organizing principle of the brain: many cortical regions respond preferentially to particular types of information or computations (Kanwisher, 2010; Zeki, 1978). However, functionally specific areas only loosely map to specific anatomical locations, limiting the ability to infer function from anatomy alone (Amunts et al., 1999; Assem et al., 2020; Fedorenko & Blank, 2020; Fedorenko & Kanwisher, 2009; Fischl et al., 2008; Frost & Goebel, 2012; Glasser et al., 2016; Tahmasebi et al., 2012; Tomaiuolo et al., 1999). Group-constrained subject-specific (GcSS) functional localization (Fedorenko et al., 2010) aims to discover broad similarities in activation patterns across individuals while accommodating interindividual anatomical differences. This approach uses whole-brain activation maps from multiple subjects to define a set of search spaces (parcels) and then identifies subject-specific functional regions of interest (fROIs) within those search spaces. Here, we conduct a large-scale fMRI analysis to determine the efficacy of the GcSS method across diverse contrasts of interest, with the goal of establishing the minimal viable sample sizes for this analytical approach.

Classic experimental designs in cognitive neuroscience commonly introduce a contrast of interest—two conditions (or groups of conditions) that differ in an important way, such as real words vs. fake words, or face images vs. non-face images. A key endeavor is then to identify and study brain areas that respond to this contrast. One traditional approach is to derive subject-specific whole-brain contrast maps and aggregate them across subjects using the group-level random-effects (RFX) method (Friston et al., 1999). A major pitfall of the RFX approach is that across-subject averaging takes place at the voxel level: voxels at anatomically equivalent locations in a shared template space are assumed to be the ‘same’ across individuals in this approach, but in reality, often belong to different functional areas in different people, reducing the sensitivity of the analysis by blurring anatomically close but functionally distinct areas (Nieto-Castañón & Fedorenko, 2012; Saxe et al., 2006). For example, Turner et al. (2018) reported that even samples exceeding 100 subjects can yield unreliable group-level activation maps.

Region-of-interest (ROI) approaches provide an alternative to whole-brain RFX analyses. By restricting analysis to predefined brain regions, ROI-based analyses increase statistical power by reducing the need for multiple comparisons correction and allowing targeted hypothesis testing (Poldrack, 2007). A central question then becomes: how best to define ROIs?

One approach is to use anatomical ROIs, defined either using a reference atlas (e.g., Destrieux et al., 2010; Glasser et al., 2016) or using participants’ individual anatomies (e.g., Nieto-Castanon et al., 2003). This method can be effective when researchers have prior hypotheses about where an effect might be observed. Empirically, statistical inference within anatomically constrained ROIs generally requires smaller sample sizes than whole-brain RFX analyses. Klapwijk et al. (2025) concluded that moderate task effects can be reliably detected with sample sizes of N = 40-60, compared with the >100 participants often needed for reliable whole-brain RFX analyses (Turner et al., 2018). Similarly, Sadil and Lindquist (2026) found that samples of N = 40-80 were sufficient to detect activations in the ten most responsive anatomical ROIs, with most contrasts showing moderate to good cross-sample consistency at these sample sizes. However, these recommended sample sizes still exceed those commonly used in fMRI studies (median N ≈ 30; Poldrack et al., 2017).

Moreover, anatomical ROIs suffer from the same problem as voxel-based whole-brain averaging approaches: they assume a one-to-one mapping between anatomy and function, which does not hold in practice. In fact, a single anatomical region commonly houses multiple functionally specialized areas (e.g., Gordon et al., 2017, 2026; Ladwig et al., 2026; Yeo et al., 2011). For example, a region in the inferior frontal cortex commonly referred to as “Broca’s area” contains regions belonging to both the frontotemporal language network and the frontoparietal multiple-demand network (Fedorenko & Blank, 2020), as well as speech planning areas (Fedorenko et al., 2024; Wolna et al., in preparation). Even fine-grained multimodal atlases (e.g., Glasser et al., 2016) benefit from subject-specific refinement because functional boundaries vary across individuals in ways that are not fully captured by a common atlas (Wang et al., 2015; Wolna et al., 2026).

To overcome the limitations of anatomy-based ROIs, cognitive neuroscience today is shifting toward precision brain imaging (Gordon et al., 2017; Gratton & Braga, 2025). Defined broadly, precision brain imaging localizes ROIs in individual brains using functional brain data (often, but not necessarily, starting with an anatomical prior). One tradition defines large-scale functional networks based on seed-based or whole-brain functional connectivity patterns (e.g., Braga & Buckner, 2017; Du et al., 2024; Shain & Fedorenko, 2025; Yeo et al., 2011). The other tradition defines functional regions of interest (fROIs) based on localizer tasks (Downing et al., 2001; Epstein & Kanwisher, 1998; Kanwisher et al., 1997; Saxe et al., 2006). The two traditions are complementary, and have generally converted to the same functionally specialized brain networks (Braga et al., 2020; DiNicola et al., 2020; Shain & Fedorenko, 2025). In this work, we focus on the fROI approach.

The group-constrained subject-specific (GcSS) method (Fedorenko et al., 2010; **Figure 1**) offers a way to define individual-level fROIs while leveraging the observation that most functional areas are loosely anchored to anatomy (Brett et al., 2002). This method first identifies group-level parcels, using a probabilistic group-level overlap map, that capture the approximate location and extent of each functional area, after which subject-specific fROIs are defined within those parcels. The GcSS method has demonstrated improved sensitivity and functional resolution relative to group-level RFX approaches (Nieto-Castañón & Fedorenko, 2012) and has been successfully applied to define networks supporting diverse functions, including vision (Julian et al., 2012; Jung et al., 2026), speech and language (Basilakos et al., 2018; Fedorenko et al., 2010; Overath et al., 2015; Regev et al., 2024), domain-general executive control (Fedorenko et al., 2013), semantic reasoning (Ivanova et al., 2025), formal reasoning (Kean et al., 2025), and intuitive physical reasoning (Fischer et al., 2016; Pramod et al., 2025).

**Figure 1.**
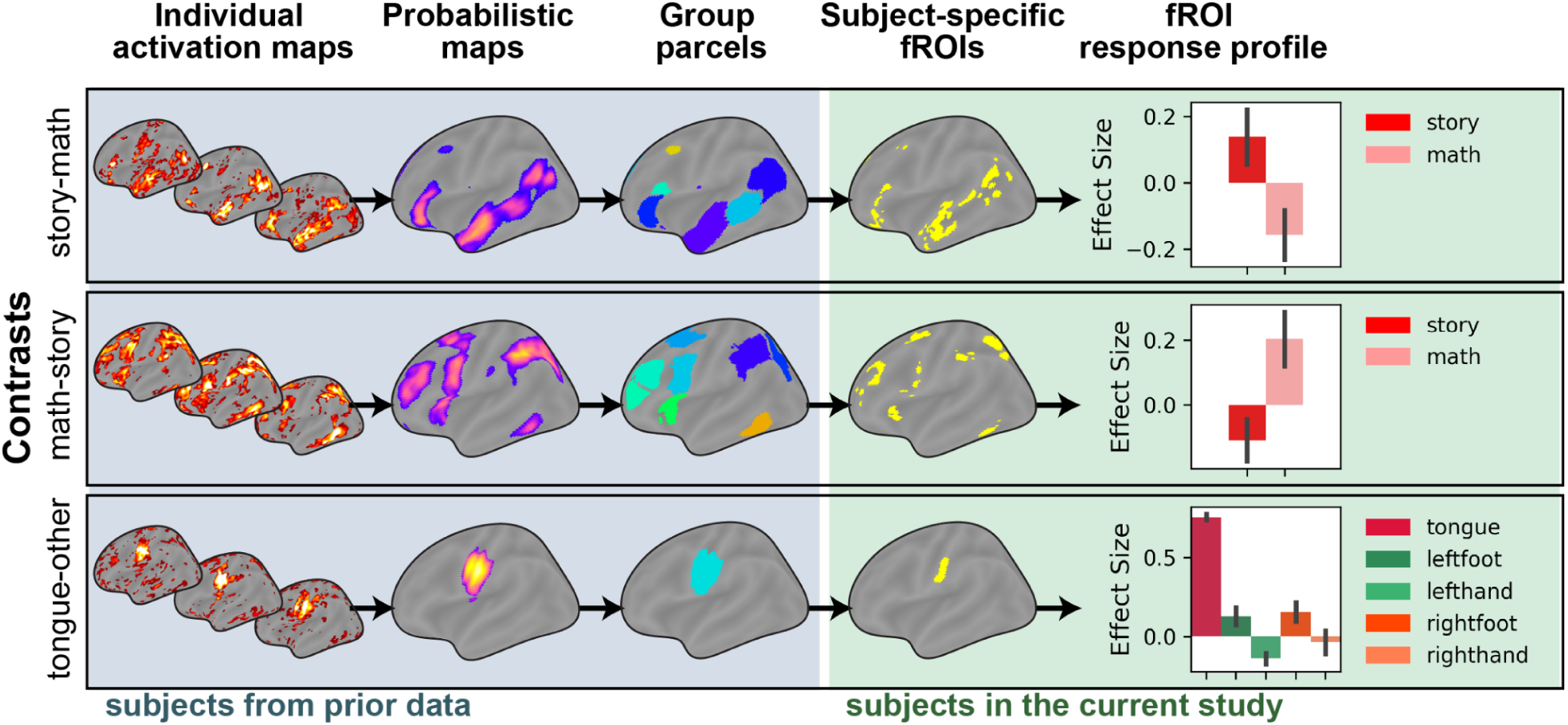
Analytical framework and examples of the group-constrained subject-specific (GcSS) method. Individual t-statistic maps for a contrast of interest are thresholded and aggregated into a probabilistic overlap map (showing the ratio of participants that had significant contrast values for each voxel). A probabilistic map is then smoothed, thresholded, and passed through a watershed algorithm to yield discrete group-level parcels. The parcels mark anatomical search spaces to define functional regions of interest (fROIs) in each subject using subject-specific localizer data. Once defined, subject-specific fROIs can be examined by measuring their response profiles in held-out data.

However, little is known about the sample sizes required for reliable GcSS analyses. Existing sample size recommendations have focused primarily on whole-brain RFX analyses or anatomically defined ROIs, leaving unanswered how many participants are needed to construct stable probabilistic maps and parcels and define reliable subject-specific fROIs to obtain robust estimates of effect sizes using the GcSS framework. Here, we ask: what sample size is required for reliable GcSS fROI-based analyses?

To address these questions, we analyzed task-based fMRI data from 900+ subjects from the Human Connectome Project (HCP; Van Essen et al., 2013), leveraging sixteen contrasts designed to localize both distributed large-scale networks (language, multiple demand, default mode, and social cognition) and more focal functionally selective brain areas (motor and visual areas). We found that reliable probabilistic overlap maps and parcels require sample sizes of ∼100 subjects; however, once the parcels are defined, robust effect size estimates for subject-specific fROIs can be obtained even with relatively small samples (N = 10-20). We also show that these scaling relationships extend both to cerebellar GcSS analyses and to surface-based cortical GcSS analyses, indicating the robustness and generalizability of the GcSS framework.

We conduct our analyses using the recently developed funROI toolbox (Gao & Ivanova, 2025). The code for this work is available at https://github.com/GT-LIT-Lab/GcSS-Scaling-Laws.

## Results

The GcSS pipeline consists of several stages (**Figure 1**). First, an initial set of contrast maps is aggregated into a probabilistic overlap map, which quantifies which portion of subjects have shown significant contrast values at each voxel. Then this map is transformed into a set of parcels. The parcels are then combined with a new subject’s contrast map to define that subject’s fROIs.

In this work, we examine sixteen contrasts spanning four tasks from the Human Connectome Project (HCP): Language, Social, Motor, and Working Memory (WM) (**Table 1**). We first verify that GcSS achieves greater effect size sensitivity than alternative functional localization methods and then characterize the scaling laws governing the estimation of (1) probabilistic maps, (2) group-level parcels, and (3) fROI effect sizes. These scaling relationships hold both in the cerebral cortex and in the cerebellum. Finally, we show that fROIs defined with the same contrast may show different response profiles; thus, although a single contrast does yield reliable fROI definitions, additional conditions can help further differentiate fROIs.

**Table 1.** Contrasts used for functional localization with GcSS. Language, social, four motor, and four visual perception regions each have a unique contrast assigned to them. The multiple demand network is hypothesized to be identifiable with two contrasts (Language math-story, Working memory 2back-0back); the default mode network is hypothesized to be identifiable with three contrasts (Working memory 0back-2back, baseline-0back, baseline-2back). The fMRI data are from the HCP-YA dataset.

| Task | Cognitive function/network | Contrast |
| --- | --- | --- |
| Language | language | story-math |
|  | multiple demand | math-story |
| Social | social | social-random |
| Motor | motor | tongue-other, leftfoot-other, lefthand-other, rightfoot-other, righthand-other |
| Working memory | multiple demand | 2back-0back |
|  | default mode | 0back-2back, baseline-2back, baseline-0back |
|  | visual perception | place-other, face-other, body-other, tool-other |

### Precision brain imaging via GcSS outperforms traditional group analyses

Our motivation for using GcSS stems from past observations that functional activation profiles are anatomically heterogeneous across subjects (see **Introduction**). Before proceeding further, we wanted to experimentally test this observation in the HCP dataset, as well as establish whether GcSS indeed provides better sensitivity than group-based ROI approaches.

To estimate between-subject variability, we compared the spatial consistency of the whole-brain contrast maps within subjects (defined using two separate runs) and across subjects. For each contrast, we found that the Dice overlap coefficient for within-subject maps was significantly higher than for between-subject maps (corrected ps < .0001; **Figure 2A**). The whole-brain activation patterns were significantly positively correlated both within and between subjects. They also showed significantly greater spatial similarity within subjects than between subjects (corrected ps < .0001, **Supplementary Figure 1**). Thus, we confirmed the existence of stable within-subject spatial patterns that are not fully captured by between-subject comparisons.

**Figure 2.**
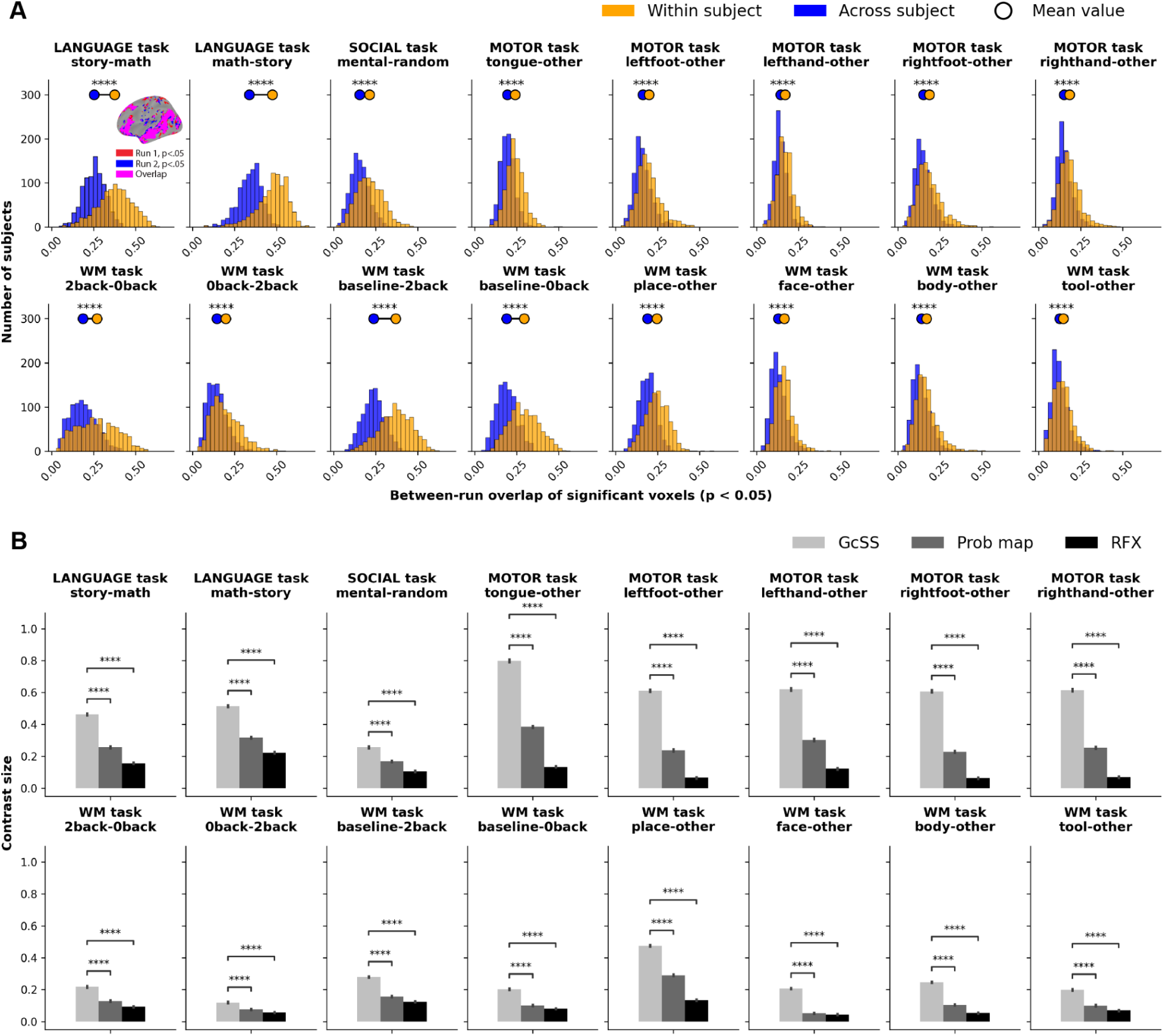
The need for subject-specific localization. (A) Dice overlap for the same contrast map from two different runs, within-subject (both runs are from the same subject) vs. across subjects (run 1 from the first subject is paired with run 2 from a different subject). (B) Effect size for the contrast of interest in ROIs defined using GcSS vs. a probabilistic group-level map vs. the traditional RFX analysis. Here and elsewhere, significance stars indicate p-value thresholds: * p < 0.05, ** p < 0.01, and *** p < 0.001. Statistical significance was assessed using one-sided Mann-Whitney U tests with Bonferroni correction for multiple comparisons.

We then tested whether GcSS, a method that aims to account for subject-specific activation patterns, results in higher effect size estimates for the contrast of interest. Using the first 450 subjects to establish parcels, probabilistic maps, or group maps defined with RFX and evaluating these on the remaining subjects (N > 450; see **Methods** for details), we found that to be the case: the effect size for each contrast of interest was significantly higher when estimated using GcSS ROIs compared to the probabilistic map (ps < .05) and RFX approaches (ps < .0001), both of which yield the same ROI masks for each subject (**Figure 2B**; see Lipkin et al, 2022, for convergent results using a different dataset). Therefore, functional localization using GcSS had significantly higher sensitivity than alternatives.

After this initial validation, we proceeded to establish the scaling laws for robust GcSS analyses.

### Scaling laws for the probabilistic overlap maps

We first examined the stability of the probabilistic overlap maps, which serve as a foundation for the parcels and—in the absence of subject-specific localizer data in a new group of subjects—as an anatomical proxy for fROIs (Lipkin et al., 2022). To track how the stability of probabilistic maps scales with sample size, we compared probabilistic maps defined in independent samples of data, with sample sizes ranging from N = 10 to N = 450. We used voxelwise intraclass correlation (ICC) as a measure of cross-sample agreement on absolute voxelwise probability estimates. For all contrasts except 0back>2back, ICCs reached good reliability (ICC > .75; Koo & Li, 2016) at N = 100; more than half (9 out of 16) reached excellent reliability (ICC > .90) by N = 200 (**Figure 3A**). All contrasts reached excellent reliability at N = 450 (the maximum sample size we examined).

**Figure 3.**
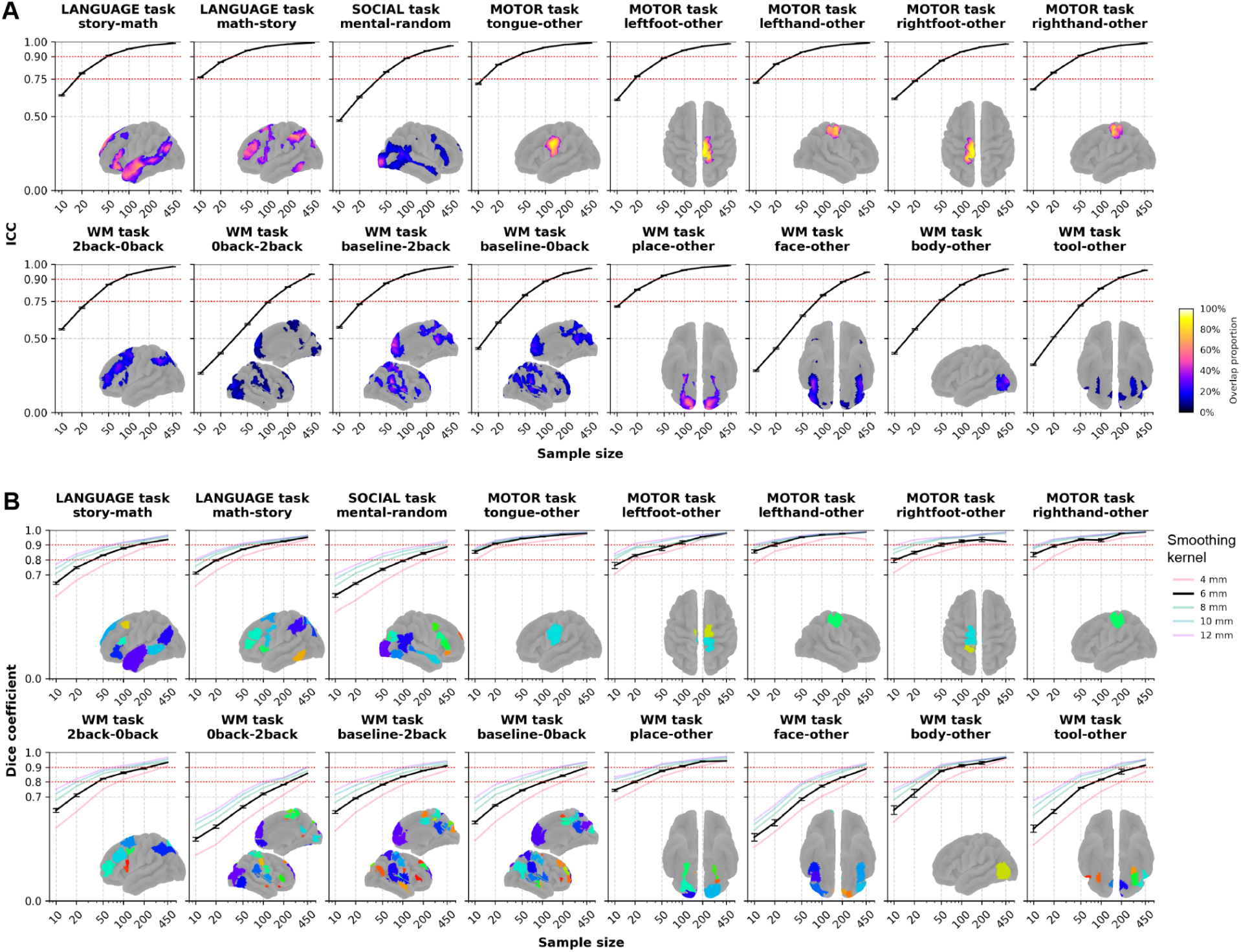
Reliability of probabilistic maps and group parcels as a function of sample size. (A) Intraclass correlation of probabilistic map estimation over voxels as a function of sample size. Error bars denote 95% confidence intervals. (B) Dice overlap coefficient of group parcels as a function of sample size. Error bars denote the standard error of the mean. Probabilistic overlap maps and group parcels were derived from the maximum sample size (N = 450).

The scaling patterns differed somewhat across contrasts. Contrasts targeting the language network (story>math), the multiple-demand network (math>story, 2back>0back), and motor regions already showed good reliability at N = 50 and excellent reliability at N = 100. The social reasoning network (social > random) was slightly less reliable, but still achieved good reliability at N = 50 and almost excellent reliability at N = 100. Most visual contrasts (face-other, body-other, and tool-other) were less reliable, at least in this dataset; however, even for them, reliability improved with increasing sample size and exceeded ICC = .75 at N = 100.

Different contrasts used to define the same functional network also had varying efficacy: the baseline>2back contrast in the working memory task exhibited higher reliability (good at N = 50) than either 0back>2back or baseline>0back, despite all three producing similar topological patterns in default mode network (DMN) regions. Similarly, the topological patterns of the multiple-demand (MD) network defined by the math > story and 2-back > 0-back contrasts were highly similar. One notable difference was that the math > story contrast consistently identified activation in the posterior inferior temporal cortex, reproducing previous observations reported by Assem et al. (2020).

### Scaling laws for the group parcels

Once the probabilistic map is defined, it is smoothed and segmented into binary anatomical parcels, which later serve as search spaces for subject-specific fROIs. To investigate how the cross-sample stability of group parcels scales with sample size, we quantified the Dice coefficient (DC) between group parcels derived from independent samples. For most contrasts, except DMN contrasts and the face perception contrast, a sufficiently large sample (N = 100) yielded substantial agreement between samples (DC ≈ 0.8) when using moderate spatial smoothing (6 mm FWHM; **Figure 3B**). With very large sample sizes (N = 450), most contrasts achieved near-perfect overlap (DC ≈ 0.9). In contrast, the DMN contrasts and the face perception contrast required larger samples (N = 200) to reach DC ≈ 0.8. The DMN network defined by the 0-back > 2-back contrast showed the lowest stability, achieving DC ≈ 0.8 only at the maximum sample size tested (N = 450).

The degree of smoothing somewhat modulated the cross-sample parcel overlap (with larger smoothing resulting in higher overlap), but did not substantially alter the scaling patterns. For noisier probabilistic maps, smaller sample sizes generally benefited more from increased smoothing. However, gains in reliability diminished with increasing smoothing magnitude and plateaued at approximately 12 mm FWHM. Importantly, even extensive smoothing could not produce substantial agreement when the sample size was very small (N = 10). Moreover, smoothing must be applied cautiously: larger kernels can distort probability distributions and shift activation peaks (Alakörkkö et al., 2017). Increasing the smoothing kernel up to 12 mm led to a progressive expansion of the search space and overly inclusive fROIs, and was associated with reduced sensitivity (**Supplementary Figure 2**).

Traditional GcSS approach also implements parcel filtering based on parcel size and the proportion of subjects showing significant voxels within that parcel. We here implemented minimal filtering based on parcel size, with the threshold of 100 voxels (corresponding to < 0.09% of the cortex and < 0.45% of the cerebellum). Because fROIs were defined as the top 10% of each parcel, this threshold ensures that at least 10 voxels are available for fROI definition. Additional filtering based on the proportion of subjects showing significant voxels within a parcel or parcel size did not meaningfully improve robustness (**Supplementary Figure 3**).

Finally, we asked whether the stability of probabilistic maps and the resulting parcels could be predicted from analyses of smaller datasets. Cross-sample consistency of probabilistic maps, estimated by comparing maps derived from independent subsets of participants, strongly predicted the sample size ultimately required to obtain stable probabilistic maps and parcels (**Supplementary Figure 4**). In contrast, neither within-subject nor between-subject between-run consistency predicted the sample size required for reliable group-level maps (**Supplementary Figure 5**).

### Scaling laws for fROI effect sizes

Next, we examined the sample size required for reliable effect size estimation in fROIs defined by the contrasts of interest using conditions from either the same or different tasks (16 contrasts of interest evaluated across a total of 17 conditions). The first 450 subjects were used to establish the group parcels. We then assessed whether the 95% confidence intervals (CIs) derived from each sample captured the ground-truth effect sizes, approximated as the effect sizes computed from all held-out subjects (N > 450). Across all sample sizes examined, more than 95% of samples contained the ground-truth effect size within their 95% CIs (**Figure 4A**).

**Figure 4.**
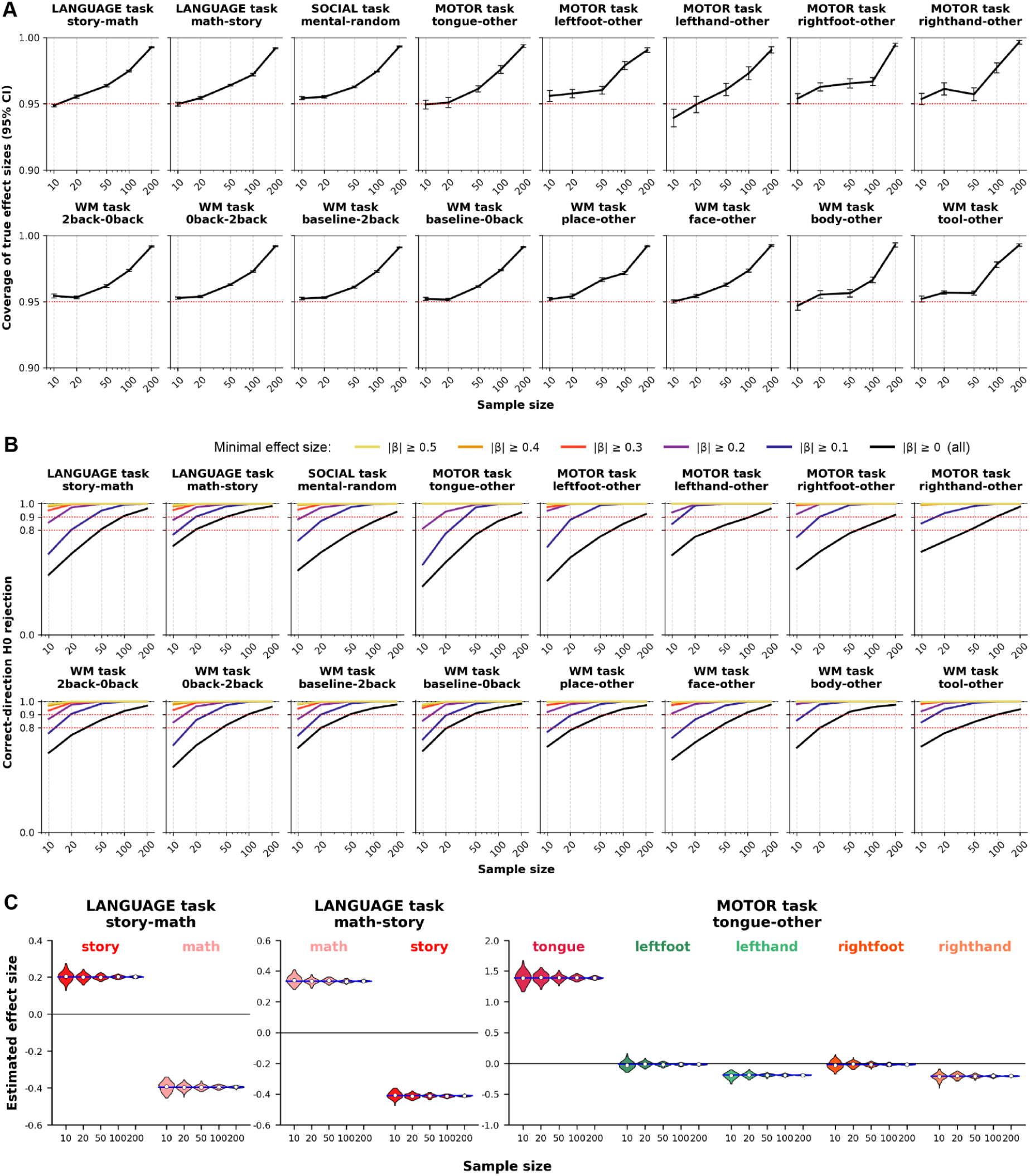
Even small sample sizes are sufficient to accurately estimate fROI effect sizes. (A) Mean proportion of samples in which the 95% CI includes the ground-truth effect size (computed from all held-out subjects, N >= 450), as a function of sample size. Error bars denote standard errors across fROIs and effect conditions. (B) Mean proportion of samples which successfully rejected the null hypothesis for significant effects, as a function of sample size and effect size. (C) Example sample size estimation by sample size. The blue lines indicate the ground-truth effect sizes. Violin plots show the distribution of estimated effect sizes across 1000 samples for each sample size.

We next evaluated sensitivity to detect effects of practical importance. We calculated the proportion of effects that lay outside 95% CI across different sample sizes. Restricting analyses to effects whose reference effect size exceeded 0.2, even very small samples (N = 10) detected these effects in 80% of cases, increasing to 90% for N = 20 (**Figure 4B**). In addition, for all sample sizes, the samples correctly retain zero in the confidence interval more than 90% of the time when the effect is non-significant. As expected, sensitivity increases with effect size, and this pattern is the same both for effect sizes derived from the same task used to define the fROI contrast and for effect sizes for conditions from a different task (**Supplementary Figure 6**). Together, these findings indicate that once the search space is robustly defined using probabilistic-map-derived parcels, fROI effect sizes can be reliably estimated using small sample sizes.

In addition to the scaling analyses, we examined the robustness of fROI sensitivity to variations in fROI definition. First, we investigated how sensitivity changes as a function of the fROI threshold. Across all contrasts of interest, both within-subject fROI definition overlap and sensitivity varied gradually with threshold, exhibiting smooth monotonic trends without any clear breakpoint or discontinuity (**Supplementary Figure 7**), suggesting gradient fROI boundaries. At the 10% threshold used in the main analyses, all fROIs exhibited substantially higher within-subject overlap than between-subject overlap, supporting the validity of this parameter choice.

We next examined the effect of the spatial search space used for fROI definition. To test the methodological value of constraining the search space, we compared fROIs defined within parcels associated with the target contrast (“correct parcels”) with fROIs defined across the whole brain (no parcels). Further, to test whether the benefit of GcSS arises simply from imposing any anatomical constraint—or instead depends on using functionally appropriate parcels—we compared fROIs defined within the “correct” parcels with those defined within parcels associated with the opposite contrast (“wrong parcels”). fROIs defined within parcels associated with the target contrast exhibited the strongest responses to that contrast (**Figure 5**). Whole-brain-defined fROIs also showed reliable effects, although with reduced effect sizes. In contrast, fROIs defined within parcels associated with the opposite contrast did not reliably yield significant responses among fROIs. Together, these results demonstrate that functionally appropriate parcel constraints improve both the sensitivity and specificity of fROI analyses.

**Figure 5.**
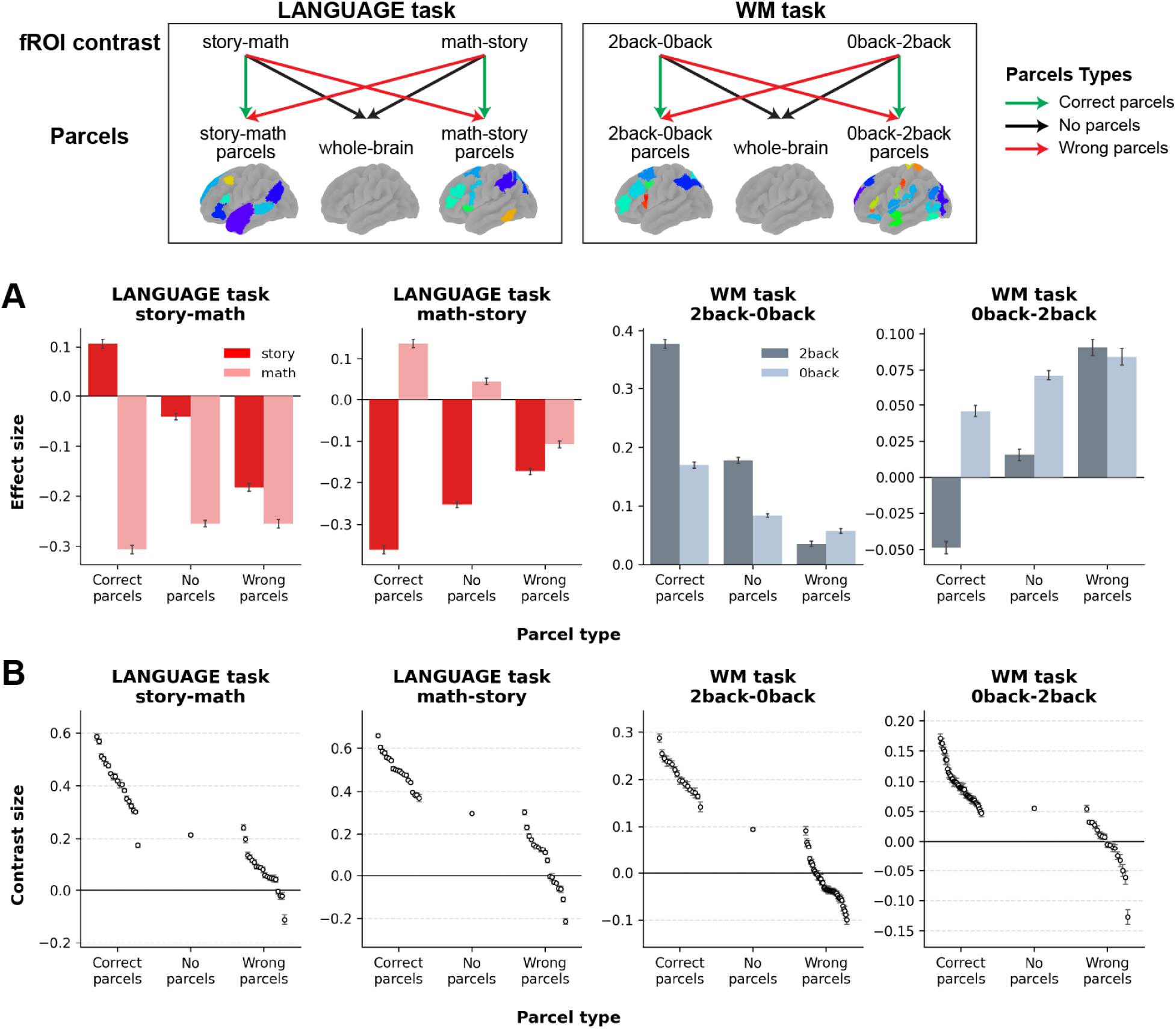
Constraining the search space with GcSS parcels has a strong influence on effect sizes in resulting fROIs. (A) Estimated effect sizes obtained when fROIs were defined by selecting significant voxels (p < .05) within the correct parcels, across the whole brain (no parcels), or using parcels defined with the opposite contrast (wrong parcels). (B) Contrast sizes by fROI, with one dot per fROI showing the across-subject mean and error bars denoting standard errors. In the no-parcels (whole-brain) condition, a single dot represents the set of all selected voxels across the brain.

### Cerebellar results mirror cortical findings

For the cerebellum, we similarly found that the GcSS method led to significantly higher contrast sizes compared with methods that use group ROIs (**Supplementary Figure 8**; corrected ps < .05). The scaling behaviors observed for probabilistic map estimation, group parcels, and effect sizes followed the same overall trend (see example contrasts in **Figure 6**). Detailed cerebellar results for all contrasts are provided in **Supplementary Figures 9** and **10**.

**Figure 6.**
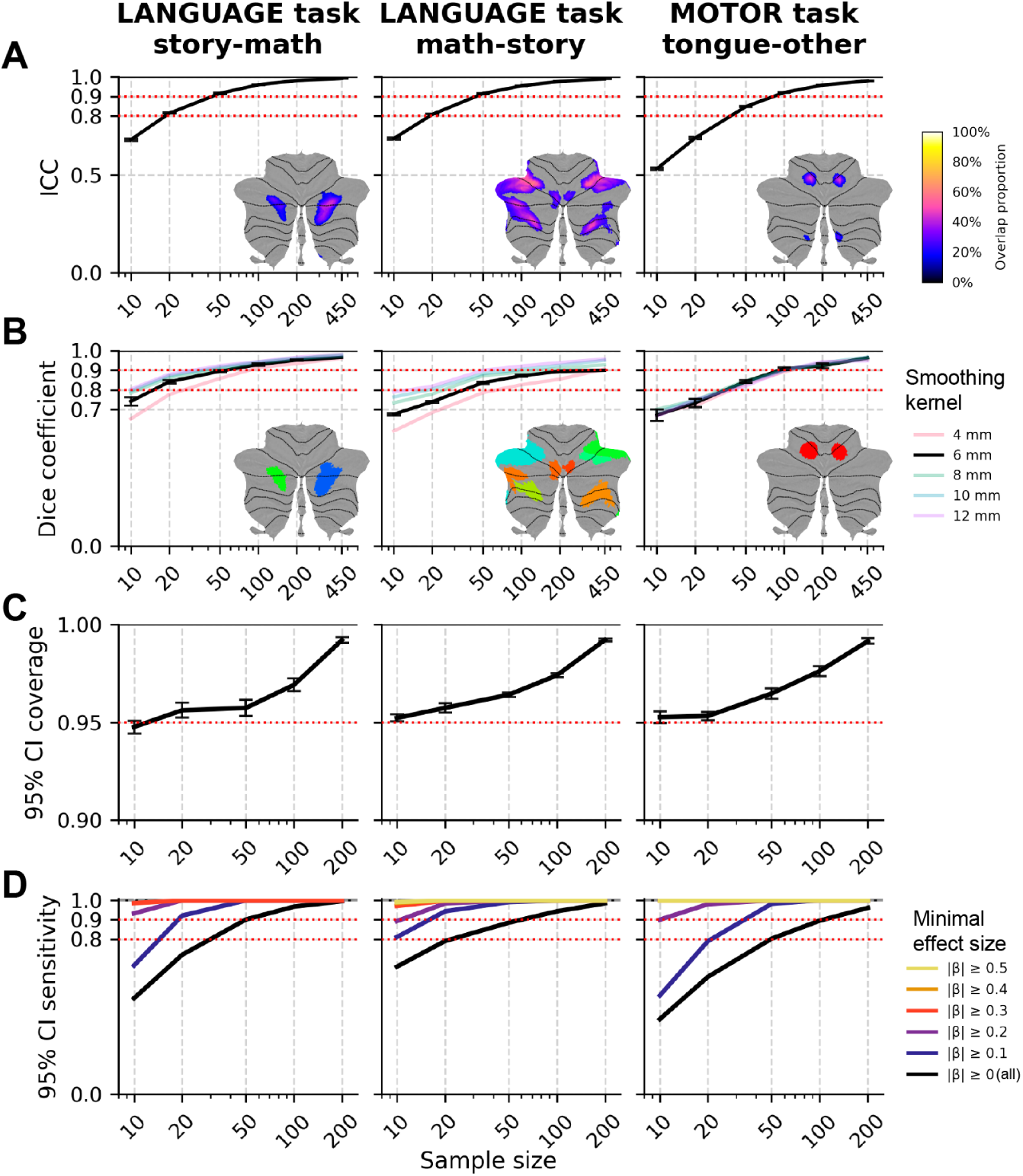
Example cerebellar results. (A) Intraclass correlation of probabilistic map estimation over voxels as a function of sample size. (B) Dice overlap coefficient of group parcels as a function of sample size. (C) Mean proportion of samples in which the 95% CI includes the ground-truth effect size, as a function of sample size. (D) Mean proportion of samples which successfully rejected the null hypothesis for significant effects, as a function of sample size.

### Surface-based analyses mirror volume-based findings

To evaluate whether our conclusions generalize beyond volume-based cortical analyses, we repeated the scaling analyses using surface-based GcSS localization. Surface-based probabilistic maps exhibited scaling behavior similar to that observed in volume space, and effect-size estimates within subject-specific fROIs were likewise comparable across sample sizes (**Supplementary Note 1**; **Supplementary Fig. 12A–B**). Thus, the scaling relationships reported here generalize across both volume and surface-based analyses.

### GcSS parcels converge across datasets

To evaluate whether GcSS-derived parcels generalize across datasets and experimental designs, we compared each set of parcels to previously published GcSS networks derived from independent datasets. **Supplementary Figure 11A** provides a qualitative visualization of parcel overlap across the brain (Dufour et al., 2013; Fedorenko et al., 2010, 2013; Julian et al., 2012). These reference atlases were derived from independent datasets using related but different contrasts of interest; they can therefore serve as an independent benchmark for assessing reproducibility.

Across all contrasts, overlap with the hypothesized target network was substantially greater than overlap with non-target networks (mean target overlap = 41.2%, mean non-target overlap = 10.6%; paired Wilcoxon test, p < 0.001; **Supplementary Figure 11B**). Furthermore, 8 of the 11 contrasts exhibited the greatest overlap with their predicted target network relative to all other reference networks, indicating a high degree of convergence across datasets and task paradigms.

Three contrasts showed notable deviations from this pattern. The SOCIAL task contrast (social > random) exhibited greater overlap with the language network than with the ToM network. The WM contrast (baseline > 0-back) showed slightly greater overlap with the ToM network than with the DMN, although the reference ToM and DMN parcels themselves exhibited substantial spatial overlap (**Supplementary Figure 11C**). Finally, the face > other contrast produced nearly equivalent overlap with face-selective visual regions and with ToM. Together, these findings indicate that GcSS parcels are largely reproducible across datasets while also highlighting cases in which task design influences the functional specificity of the resulting parcels.

### fROIs defined with the same contrast can vary in their response profiles

All 16 examined contrasts could successfully be used to define reliable probabilistic maps, group parcels, and subject-specific fROIs. How should we interpret the resulting fROIs? Are fROIs identified with the contrast A>B selective for A over all other conditions? And do all fROIs identified with the same contrast exhibit the same functional response profile?

To address these questions, we examined the functional response profiles of our fROIs more closely. We expected contrasts targeting domain-general systems to identify regions exhibiting broad response profiles across multiple conditions, with similar response profiles across regions defined using different contrasts targeting the same network. In contrast, we expected contrasts targeting domain-specific systems to identify regions with more selective response profiles tied to particular conditions of interest.

Consistent with this expectation (**Figure 7A**), fROIs localized using contrasts targeting the domain-general MD network exhibited highly similar response profiles when defined using two different contrasts. Response profiles of fROIs defined using math>story and 2back>0back were nearly identical (r = 0.99, p < .001). Similarly, for the default network, response profiles were strongly correlated across fROIs defined using the 0back>2back, baseline>2back, and baseline>0back contrasts (r = 0.77–0.98, ps < .001). Together, these results indicate that contrasts targeting the same domain-general network tend to identify fROIs with highly similar functional response profiles.

**Figure 7.**
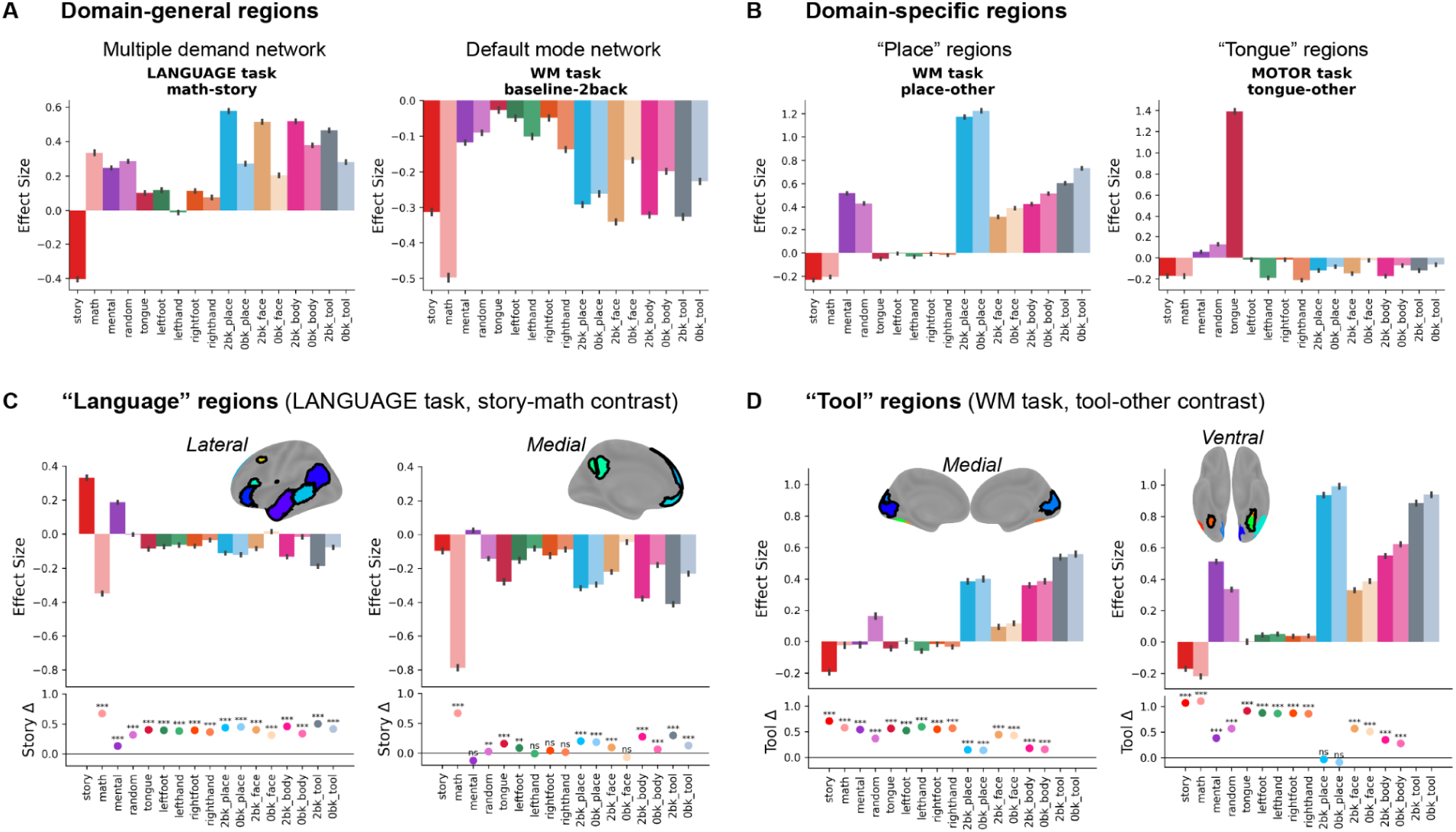
fROIs defined by a single contrast can exhibit varied functional response profiles. **(A)** Domain-general regions have broad response profiles. (B) Domain-specific regions have selective response profiles. (C) Lateral and medial fROIs defined with the story-math contrast show distinct response profiles (top) and target contrast preferences (bottom), with only the lateral fROIs showing language selectivity. (D) Tool regions show only weak (left) or absent (right) preference for tools over places. Target contrast preferences were measured by comparing the average response in the reference condition (story for language fROIs; tool for tool fROIs) with the average response in each of the remaining conditions.

For most other networks (**Figure 7B**), when responses were averaged across all fROIs within each network, nearly all contrasts used to localize functionally selective networks/regions exhibited the expected functional selectivity. Specifically, for contrasts targeting all networks except the MD and DMN, the target conditions elicited significantly stronger responses (ps < .001) than other conditions from the same or different tasks. For instance, motor fROIs defined using the tongue>other contrast responded most strongly to the tongue condition when comparing responses across all 17 conditions. Similarly, place fROIs defined using the place>other contrast responded most strongly during visual perception of place stimuli among all conditions.

There were, however, some exceptions to this overall pattern. First, the language network did not show significantly stronger responses to the story condition than to the mental condition in the social cognition task when we considered its average response across fROIs. Upon further inspection, individual fROIs showed substantial heterogeneity (**Figure 7C**). Specifically, left lateral language regions showed strong selectivity for the story condition (ps < .001), whereas medial regions exhibited deactivation during cognitively demanding tasks rather than selective activation for stories, suggesting that the story>math contrast identified two different response profiles.

Second, tool perception regions, on average, did not show significantly greater activation for the 2-back tools condition than for the 0-back places condition. Here, too, individual fROIs exhibited heterogeneous response profiles (**Figure 7D**). Medial regions showed selectivity for tools (ps < .001), whereas ventral regions were strongly driven by place-related activation, with responses to tools not significantly exceeding those to places.

Lastly, we observed that body perception regions (extrastriate body area; EBA) did not show stronger responses to body perception conditions (0-back/2-back) than to the mental or random conditions in the social-cognition task, consistent with prior findings that EBA is responsive to social interaction (Landsiedel et al., 2022).

## Discussion

One key goal of cognitive neuroscience is to discover the functional architecture of the human brain. Achieving this goal requires taking into account the now well-established inter-individual topographic variability. GcSS analyses (Fedorenko et al., 2010; Julian et al., 2012) provide a principled way for identifying subject-specific regions of interest (ROIs) that have the same functional response profile. Using the HCP-YA dataset, we derived the scaling curves for different stages of the GcSS pipeline. We found that the initial steps—defining group-level search spaces (parcels)—yield stable results for sample sizes of N=100 or up, with some variability depending on the specific contrast. Once the parcels are defined, however, they can be used to define subject-specific fROIs and accurately estimate population-level effect sizes using a sample as small as N=10. We therefore show how group priors derived from large-scale fMRI datasets make it possible to conduct subsequent fMRI studies—critical for testing a multitude of hypotheses about the function of the relevant brain areas—with much smaller samples, as long as those follow-up studies define subject-specific ROIs using functional data.

The reason to use the GcSS approach over fixed-ROI methods (where the same set of voxels is used for each participant) is the substantial inter-individual variability in the precise locations of functionally specialized networks and regions (Nieto-Castañón & Fedorenko, 2012). In contrast, within-individual activation patterns exhibit strong spatial consistency across sessions (Gordon et al., 2017; Laumann et al., 2015) and remain stable over long time periods (Poldrack et al., 2015). Consistent with prior work (e.g., Gao et al., 2026; Gordon et al., 2017; Tavor et al., 2016), we observed markedly stronger within-subject than between-subject consistency in activation locations and patterns. Moreover, in line with prior work (Lipkin et al., 2022; Nieto-Castañón & Fedorenko, 2012; Wolna et al., 2026), we show that the GcSS method for functional localization provides greater sensitivity than traditional group-level RFX approaches. We observed this advantage not only for contrasts aimed to identify cognitive networks (language, MD, DMN) but also higher-order visual areas (faces, places, bodies) and even motor areas, where anatomy and function align relatively closely (e.g., Hackett, 2015; Hinds et al., 2009)—although the within-subject vs. between-subject divide for the motor areas was smaller. These findings support the use of subject-specific fROI definitions, which explicitly accommodate individual variability in functional topography.

### Sample size requirements differ for parcel definition vs. fROI effect size estimation

We found that constructing reliable group-level parcels for use as search spaces in later GcSS analyses benefits from relatively large samples (N = 100–200). Consistent with Sadil and Lindquist (2026), the sample size required for reliable estimation of probabilistic maps and group parcels varies by contrast: HCP’s language and motor contrasts achieve good-to-excellent reliability with relatively small samples (N = 50), whereas social and working memory contrasts require substantially larger samples (up to N = 200) to reach good reliability. We showed that increasing the smoothing kernel up to 12 mm can improve parcel reliability with a smaller sample size. However, because spatial smoothing degrades fROI sensitivity, we recommend using large sample sizes with mild smoothing when exploring new contrasts for fROI definition, and applying strong smoothing only when such sample sizes are not practically attainable.

Our key finding is that when using functional localization with pre-defined parcels to estimate effect sizes in individual subjects, small sample sizes are sufficient to achieve accurate population-level estimates. This result paints a much more optimistic picture about the reliability of fMRI as a method compared to prior studies based on anatomical ROIs or RFX methods (Klapwijk et al., 2025; Turner et al., 2018). Functional localization markedly reduces sample size requirements, enabling fMRI studies that stay within reasonable single-lab research budgets.

### The scaling laws generalize across GcSS implementations

The cerebellar analyses largely mirrored the scaling behavior observed in cortex. Probabilistic maps and group parcels in the cerebellum required similar sample sizes as in cortex to reach comparable reliability. These findings demonstrate that GcSS-based functional localization is applicable to cerebellar analyses (Basilakos et al., 2018; Casto et al., 2026; Ivanova et al., 2025; Wolna et al., 2026). This extension is particularly relevant given growing evidence that the cerebellum contributes not only to motor control, but also to higher-order cognitive functions including language, working memory, and social cognition (Buckner, 2013; Guell et al., 2018; Van Overwalle et al., 2014). Similarly, the scaling relationships generalized to surface-based cortical GcSS analyses, which exhibited comparable scaling behavior for probabilistic maps and similar effect-size estimates for subject-specific fROIs. Together, these findings indicate that the sample size recommendations derived here are robust across multiple implementations of the GcSS framework.

### Recommendations for using GcSS with a new contrast

Although we have provided general sample size suggestions for GcSS parcel derivation, the required sample size will depend on a particular contrast. To establish whether the GcSS approach yields stable probabilistic maps and group parcels with a new contrast, we recommend using a split-half approach that we have adopted for the analyses here: applying the GcSS procedure to two halves of the data and measuring the similarity of the resulting maps. As shown in **Supplementary Figure 4**, cross-sample probabilistic map consistency provides a useful indicator of the sample size that would be sufficient to obtain stable probabilistic maps and parcels. For smaller datasets, another possible approach is to iteratively leave out one or several participants and examine how the resulting parcels differ from those derived from the full dataset, which may provide a practical heuristic for assessing parcel stability when split-half analyses are not feasible. If all or some parcels are unstable, they may need to be excluded from the final set—or, at least, the fROI derived from these parcels should be treated with caution. A probabilistic map derived from a smaller number of participants than the one recommended might need to be thresholded at a higher inter-subject overlap value than usual, as those high-overlap values are the most likely to hold in a larger sample.

What if the contrast of interest does not have a large enough dataset to derive GcSS parcels? In that case, fROIs may be defined within other anatomical search spaces, such as anatomical ROIs selected based on past literature or masks derived through meta-analyses (e.g., ALE studies (Eickhoff et al., 2012) or aggregation platforms like Neurosynth (Yarkoni et al., 2011)). The GcSS parcels can also be derived with a contrast that’s different from the contrast used for fROI definition, as long as the contrasts are thought to target the same set of brain regions (e.g., math>story and 2back>0back in our study yielded fairly similar parcels, consistent with the multiple demand network’s reported domain generality; Assem et al., 2020; Fedorenko et al., 2013; Shashidhara et al., 2019). Finally, even GcSS parcels derived from smaller datasets can be informative, especially for a well-designed contrast; as long as the fROI response profiles are estimated in data that were not used to define them (Kriegeskorte et al., 2009), the reported results are valid. The main caveat for such low-powered parcels is a possibility of false negatives: just because a brain area did not get its own parcel, it does not mean that it was not involved.

Lastly, we show that beyond adequate sample size, the validity of fROIs depends critically on contrast design. Contrasts that differ substantially in task demands may spuriously recruit regions of the multiple demand network (if the critical condition is more demanding) or the default mode network (if the control condition is more demanding). For example, for language, current best practices use contrasts between language processing and a perceptually matched control condition, such as reading sentences versus lists of nonwords (e.g., Fedorenko et al., 2010) or listening to sentences versus backward speech (Malik-Moraleda, Ayyash et al. 2022), which are matched for task demands (Gao et al., 2026). Importantly, robust activation for a given contrast does not by itself validate the contrast as meaningful (i.e., isolating the relevant perceptual or cognitive process). When the critical and control conditions are poorly matched, the profiles of localized fROIs will reflect all the confounds inherent in the contrast.

### GcSS complements other precision neuroscience methods

GcSS relies on traditional cognitive neuroscience experimental designs: there is a paradigm with a critical contrast of interest, and the goal is to identify brain areas that are sensitive to that contrast. Alternative approaches may bypass task contrasts altogether and rely on intrinsic brain dynamics to identify functionally coherent brain areas. Such approaches typically cluster brain voxels based on their time series during rest (Kong et al., 2019; Yeo et al., 2011); and/or during task (Du et al., 2025; Shain & Fedorenko, 2025).

Some functional network parcellation methods—most notably, the multi-session hierarchical Bayesian model approach (Du et al., 2024; Kong et al., 2019)—are conceptually similar to GcSS in that they rely on a group prior, which is derived from a large set of subjects. This prior is then combined with individual-specific fMRI data to derive subject-specific networks of interest in a smaller number of subjects. However, the prior is not necessary: some approaches successfully identify subject-level networks using fully data-driven approaches such as ICA (Shain & Fedorenko, 2025), although interpreting the resulting networks still requires functional task data.

Convergent results from task-based functional localization and timeseries-based network parcellation can indicate that the task contrast loads on a specific functional brain network (e.g., the language network: Braga et al., 2020; Shain & Fedorenko, 2025). Conversely, if there is a difference between the task-evoked and timeseries-based brain maps, it may indicate that a contrast engages multiple functional networks. For example, in the HCP dataset, the ‘language’ contrast, story-math, likely recruits two different functional networks: the language network, which is sensitive to sentence-level meaning (Fedorenko et al., 2011) and the default network, which tracks story-level meanings (Braga et al., 2020; Simony et al., 2016).

In summary, our results demonstrate that GcSS functional localization provides a scalable and reliable framework for task-based fMRI analyses. Across a diverse set of contrasts spanning language, social cognition, executive function, perception, and motor control, we show that the different stages of the GcSS pipeline have distinct sample-size requirements: robust probabilistic maps and group parcels generally require samples on the order of 100–200 participants, whereas population-level effect-size estimation within subject-specific fROIs can be achieved even in studies with as few as 10-20 participants. These findings reconcile two seemingly conflicting observations in the fMRI literature: large samples are often necessary to obtain stable group-level maps, yet carefully localized subject-specific analyses can yield meaningful and reproducible results in much smaller cohorts. More broadly, our results support a precision-neuroscience approach in which large shared datasets are used to establish robust anatomical search spaces, while individual functional localization is used to test diverse hypotheses about the relevant regions by richly characterizing their responses to new conditions in new participants. By quantifying the scaling laws of GcSS analyses in both cortex and cerebellum, we provide practical guidance for future studies, a framework for designing fMRI experiments that are both statistically rigorous and economically feasible, and tools that enable these analyses.

## Methods

### Data

We used task-based fMRI data from four tasks in the HCP-Young Adult (HCP-YA) dataset (Van Essen et al., 2012): LANGUAGE, SOCIAL, MOTOR, and WM (working memory). Across these tasks, we selected 16 contrasts spanning a range of cognitive domains (see **Table 1**). All analyses were performed on the minimally preprocessed fMRI data, which were nonlinearly registered to MNI152 space and resampled to 2 mm isotropic voxel resolution as part of the HCP minimal preprocessing pipeline (Glasser et al., 2013).

Quality control (QC) procedures were applied prior to inclusion. We excluded: (1) Runs with unresolved known issues, or segmentation errors and temporal instabilities in acquisition (QC issue codes B and C; HCP S1200 Release Reference Manual, 2018); (2) Runs in which more than 10% of volumes were flagged as motion outliers (framewise displacement > 0.5 mm). For each task, we further required that subjects had two complete, QC-passed runs in order to be included in the final dataset. The final sample sizes were 919 subjects for LANGUAGE, 945 for WM, 908 for MOTOR, and 911 for SOCIAL.

### First level modeling

We estimated voxelwise responses with a Generalized Linear Model (GLM) that modeled each experimental condition using a boxcar function convolved with the Glover hemodynamic response function (Glover, 1999). We addressed temporal autocorrelations in the BOLD signal by applying a high-pass filter with cutoff frequencies of 0.005 Hz for LANGUAGE, 0.002 Hz for MOTOR, and 0.004 Hz for the remaining tasks, followed by whitening using an AR(1) model. The design matrix included first-order temporal derivatives for each condition, six motion parameters, and one-hot motion regressors for volumes exceeding 0.5 mm framewise displacement. Finally, we applied spatial smoothing using a Gaussian kernel with a 4-mm FWHM.

We conducted first-level modeling and all subsequent GcSS analyses using the open-source Python package funROI (Gao & Ivanova, 2025). Software supporting GcSS analyses is additionally available in MATLAB (spm_ss; Nieto-Castañón & Fedorenko, 2012).

### Evaluating inter-and intra-subject consistency of functional activation

For within-subject comparisons, we used the first and second runs from the same subject; for across-subject comparisons, we used the first run of one subject and the second run of the subsequent subject.

To assess not only the location of significant voxels but also the similarity of activation patterns, we computed spatial correlations of contrast-related activation values within significant voxels, again comparing within-subject to across-subject pairs. Differences in Dice overlap and spatial correlation (within-subject > across-subject) were tested using paired t-tests, with Bonferroni correction applied separately for each test type.

### Group-constrained subject specific (GcSS) analyses

We conducted GcSS analyses following the steps described in Fedorenko et al (2010), with small adjustments. For each contrast, we generated a probabilistic map by averaging voxel significance across both runs across subjects and applied a watershed algorithm to segment the map into parcels. For watershed segmentation of probabilistic maps into parcels, unless otherwise specified, we applied a 6 mm FWHM spatial smoothing and determined the probability threshold using Otsu’s method (Otsu, 1975). See **Supplementary Figures 10** and **11** for a visualization of probabilistic maps and parcels for each contrast.

For probability-map and group-parcel analyses, the number of available independent samples was 10 samples for N = 10, 20, and 50; 9 samples for N = 100; 4 samples for N = 200; and 2 samples for N = 450. We assessed the reliability of probabilistic map estimation and group parcellations by calculating the consistency intraclass correlation coefficient for single measurements (ICC(C,1)) and pairwise dice overlap coefficients, respectively, across different samples within the same sample size, and see how it scales as a function of sample size.

After parcel generation, we examined the scaling laws governing effect size estimation within subject-specific fROIs. For each contrast, fROIs were defined as the top 10% of voxels most responsive to the contrast of interest within each parcel. This threshold is conservative, smaller than the average proportion of significant voxels (p < .001) observed within parcels (see **Supplementary Table 1**), and was applied using the robust parcel definitions derived from the initial sample of 450 subjects. Using the remaining subjects, we tested the reliability of estimated effect sizes over fROIs and conditions for sample sizes of N = 10, 20 (10 samples each), N = 50 (9 samples), N = 100 (4 samples), N = 200 (2 samples). Separate runs were used for defining fROIs and estimating their effect sizes of same-task conditions to avoid circularity (double dipping; Kriegeskorte et al., 2009).

To assess validity of the effect sizes, for each localizer contrast and for each condition, we repeatedly estimated effect sizes by randomly drawing 100 samples of each sample size from the heldout subjects. To estimate the validity of confidence intervals, we then calculated, for each sample size, the proportion of samples with ground-truth effect sizes (mean effect across all held-out subjects) contained within the 95% confidence interval of the estimated effect sizes. We further assessed sensitivity and specificity of effect size estimation: Sensitivity was defined as the proportion of ground-truth significant effects exceeding a minimal effect size threshold that were correctly identified as significant in the expected direction. Specificity was defined as the proportion of ground-truth non-significant effects that were correctly identified as non-significant.

To examine the selectivity of non-domain-general networks or regions, we tested whether, in those ROIs (language: story>math; social cognition: mental>random; motor; and visual working memory), the main conditions elicited significantly higher responses than other conditions from the same or different tasks. A Bonferroni correction was applied to control for multiple comparisons within each contrast.

To assess the robustness of our findings to fROI definition choices, we conducted two sensitivity analyses.

For the threshold sensitivity analysis, the main analyses defined fROIs as the top 10% of voxels showing the strongest response to the contrast of interest within each parcel. This threshold is relatively conservative, as it is smaller than the average proportion of significant voxels (p < .001) observed within parcels (**Supplementary Table 1**). We therefore repeated the analyses across a range of fROI percentage thresholds and evaluated the resulting contrast sensitivity.

For the search-space sensitivity analysis, we compared fROIs defined within (1) parcels associated with the target contrast, (2) parcels associated with the opposite contrast, and (3) the whole brain. This analysis focused on pairs of networks whose defining contrasts are exact opposites: the language network (story > math) versus the MD network (math > story), and the DMN (0-back > 2-back) versus the MD network (2-back > 0-back). For each search space, fROIs were defined as voxels showing a significant response to the target contrast (p < .05), and contrast effect sizes were computed using the same procedures as in the main analyses.

### Comparing sensitivity across functional localization methods

To compare the sensitivity of fROIs defined by the GcSS method with those derived from fixed-fROI approaches, we examined responses to the contrast of interest across three methods: (1) subject-specific fROIs defined using the GcSS procedure, (2) group-level random-effects (RFX) fROIs, and (3) weighted probabilistic map-based fROIs. For each task, the first 450 subjects were used to derive the group-level parcels, fROIs, or probabilistic maps, and the remaining subjects were used to evaluate the contrast magnitude. In the GcSS method, subject-specific fROIs were conservatively defined as all voxels within each parcel that showed significant activation (p < .05) for that subject. Group-level RFX fROIs were defined by thresholding the second-level contrast maps at p < .05. For the probabilistic map approach, fROIs were defined by selecting the top N voxels, where N corresponded to the average number of significant voxels across subjects (p < .05). The effect of interest was computed as a weighted average of voxelwise effects, with weights given by the voxelwise probabilistic map values. Finally, we investigated the sensitivity comparison when the voxel counts of the fixed-fROI methods to the average number of voxels included in the GcSS-defined fROIs.

### Cross-dataset parcel overlap analysis

To assess the consistency of GcSS-derived parcels across datasets, we compared the parcels derived from each contrast to previously published GcSS parcels when a corresponding reference network was available. Reference networks included the language, MD, DMN, ToM, and visual perception networks (Dufour et al., 2013; Fedorenko et al., 2010, 2013; Julian et al., 2012).

For each contrast of interest, we quantified spatial overlap between the derived parcel and each reference network using Dice coefficient. We then compared overlap with the hypothesized target network against overlap with non-target networks across contrasts. Statistical significance was assessed using paired Wilcoxon signed-rank tests.

### Code Availability

The code used to perform all analyses and generate the figures in this study is available at: https://github.com/GT-LIT-Lab/GcSS-Scaling-Laws.

## Acknowledgments

Data were provided by the Human Connectome Project, WU-Minn Consortium (Principal Investigators: David Van Essen and Kamil Ugurbil; 1U54MH091657) funded by the 16 NIH Institutes and Centers that support the NIH Blueprint for Neuroscience Research; and by the McDonnell Center for Systems Neuroscience at Washington University. This project was supported by startup funding from Georgia Tech. We thank Evelina Fedorenko and Jin Li for their feedback on this work.

## Author contributions

RG: Conceptualization, Data curation, Formal analysis, Methodology, Software, Validation, Visualization, Writing – Original Draft; AI: Conceptualization, Funding acquisition, Methodology, Project administration, Resources, Supervision, Validation, Writing – Review & Editing.

## Declaration of generative AI and AI-assisted technologies in the writing process

During the preparation of this work, the authors used generative AI tools to improve language and readability. After using this tool, the authors reviewed and edited the content as needed and took full responsibility for the content of the publication.

