## Supplementary Materials for "Scaling laws for group-constrained, subject-specific task fMRI analyses"

### Supplementary Information

**Supplementary Table 1.** Average proportion of significant voxels within parcels.

| Task | Contrast | Cerebral Cortex | | | | Cerebellum | | | |
|---|---|---|---|---|---|---|---|---|---|
| | | p<.05 | p<.01 | p<.001 | p<.0001 | p<.05 | p<.01 | p<.001 | p<.0001 |
| LANGUAGE | math-story | 0.647 | 0.566 | 0.474 | 0.401 | 0.606 | 0.499 | 0.380 | 0.292 |
| | story-math | 0.570 | 0.480 | 0.384 | 0.312 | 0.527 | 0.434 | 0.336 | 0.264 |
| SOCIAL | mental-random | 0.408 | 0.279 | 0.166 | 0.101 | 0.357 | 0.223 | 0.117 | 0.063 |
| MOTOR | tongue-other | 0.781 | 0.721 | 0.650 | 0.590 | 0.466 | 0.369 | 0.271 | 0.203 |
| | leftfoot-other | 0.696 | 0.618 | 0.529 | 0.459 | 0.517 | 0.375 | 0.240 | 0.155 |
| | lefthand-other | 0.736 | 0.670 | 0.592 | 0.528 | 0.469 | 0.345 | 0.228 | 0.154 |
| | rightfoot-other | 0.702 | 0.622 | 0.531 | 0.459 | 0.509 | 0.367 | 0.234 | 0.152 |
| | righthand-other | 0.729 | 0.661 | 0.583 | 0.520 | 0.499 | 0.374 | 0.254 | 0.176 |
| WM | 2back-0back | 0.439 | 0.338 | 0.243 | 0.179 | 0.399 | 0.281 | 0.177 | 0.115 |
| | 0back-2back | 0.325 | 0.213 | 0.121 | 0.071 | 0.216 | 0.131 | 0.068 | 0.037 |
| | baseline-2back | 0.476 | 0.375 | 0.275 | 0.206 | 0.357 | 0.261 | 0.173 | 0.117 |
| | baseline-0back | 0.399 | 0.293 | 0.196 | 0.135 | 0.286 | 0.188 | 0.108 | 0.064 |
| | place-other | 0.617 | 0.516 | 0.406 | 0.322 | | | | |
| | face-other | 0.292 | 0.205 | 0.132 | 0.091 | | | | |
| | body-other | 0.530 | 0.421 | 0.311 | 0.235 | | | | |
| | tool-other | 0.385 | 0.264 | 0.159 | 0.099 | | | | |

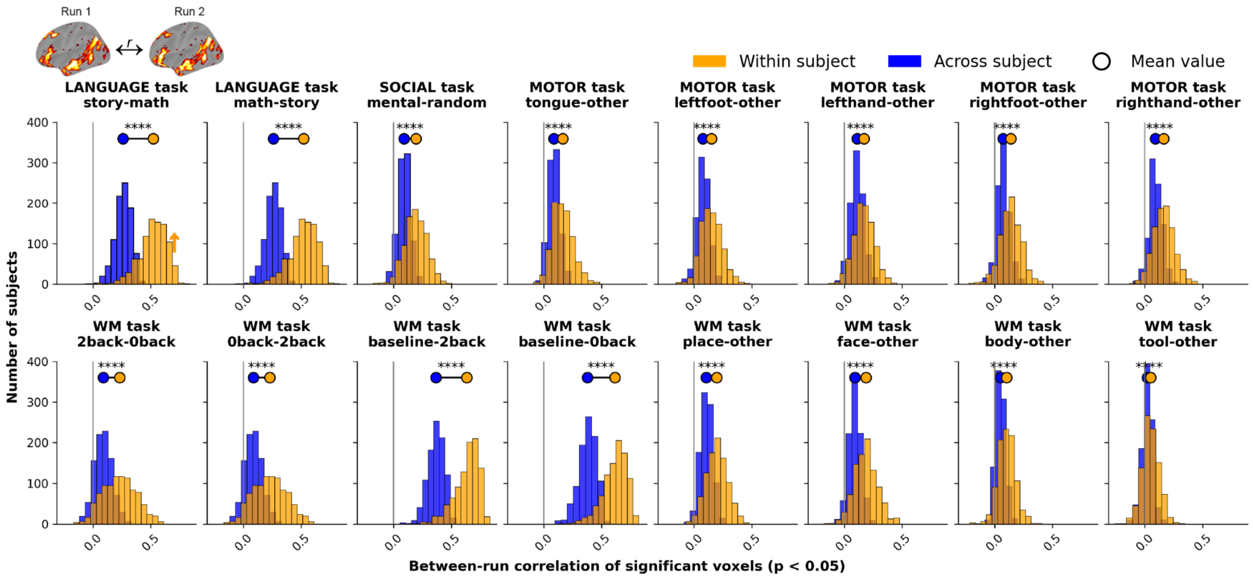


**Supplementary Figure 1.** Pearson correlation for the same contrast map from two different runs, within-subject (both runs are from the same subject) vs. across subjects (run 1 from the first subject is paired with run 2 from a different subject).

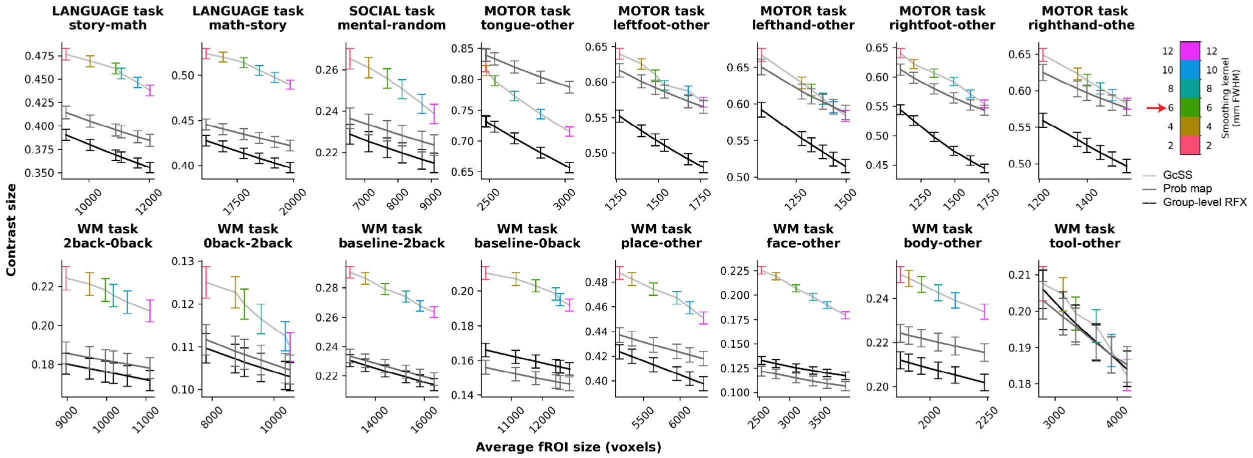


**Supplementary Figure 2.** Contrast size of fROIs defined using different methods with matched average fROI sizes.

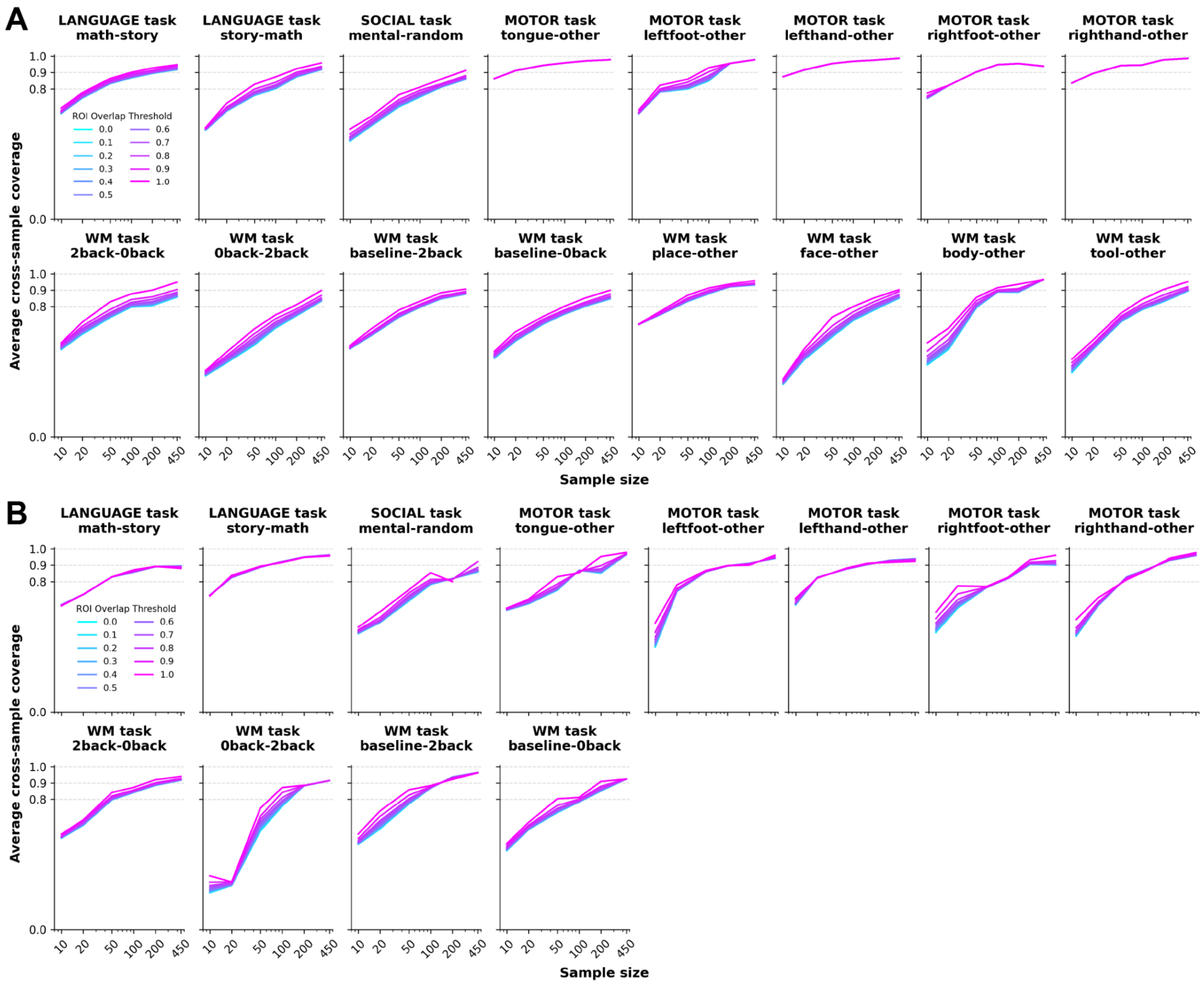


**Supplementary Figure 3.** Mean across-sample parcel coverage as a function of sample size and ROI overlap-based filtering. (A) Cortical parcels. (B) Cerebellar parcels. ROI overlap filtering yields minimal gains in coverage across sample sizes.

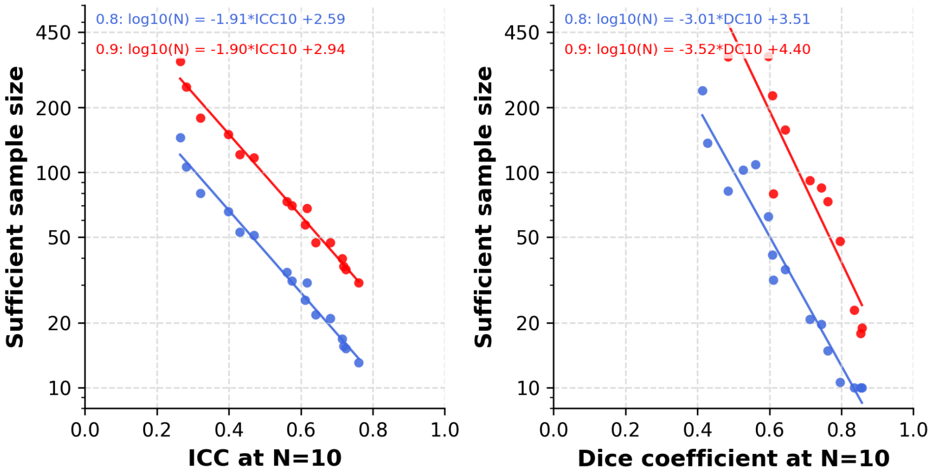


**Supplementary Figure 4.** Sufficient sample sizes based on between-sample consistency of probabilistic maps (ICC) and parcels (DC) using N = 10 samples. For each contrast, consistency was modeled as a function of log-transformed sample size, and the sufficient sample size was estimated by linear interpolation of this relationship to the predefined consistency threshold.

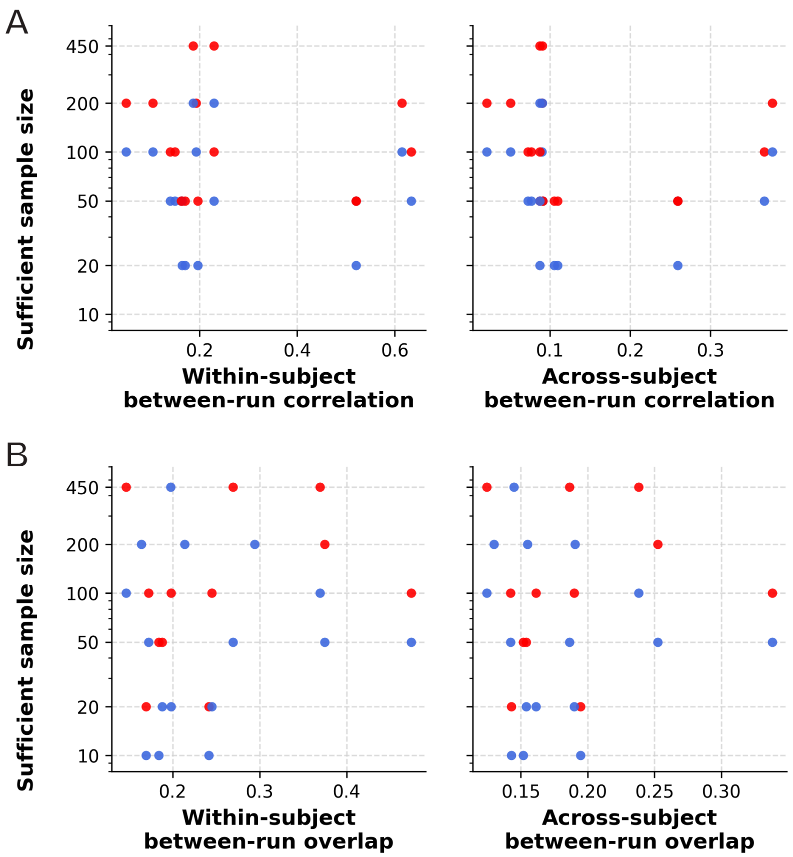


**Supplementary Figure 5.** Sufficient sample sizes by (A) between-run whole-brain correlations and (B) between-run overlap of significant voxels.

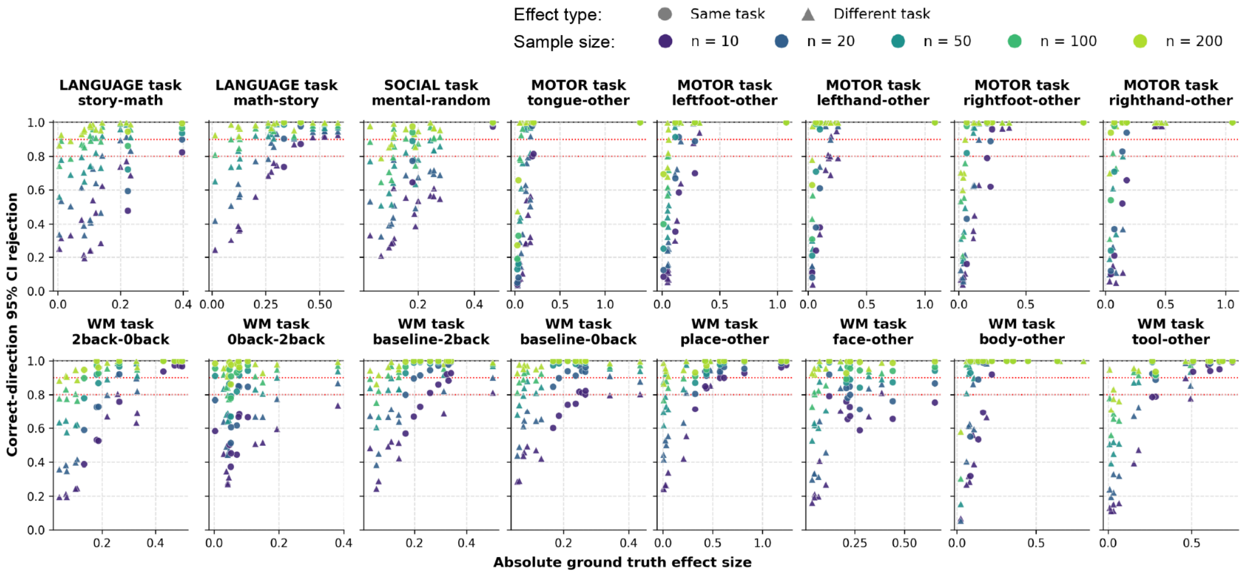


**Supplementary Figure 6.** Mean proportion of samples which successfully rejected the null hypothesis for significant effects, as a function of sample size, effect size, and effect type. Effects evaluated on the same task used to define the fROI and on a different task are indicated by different marker shapes.

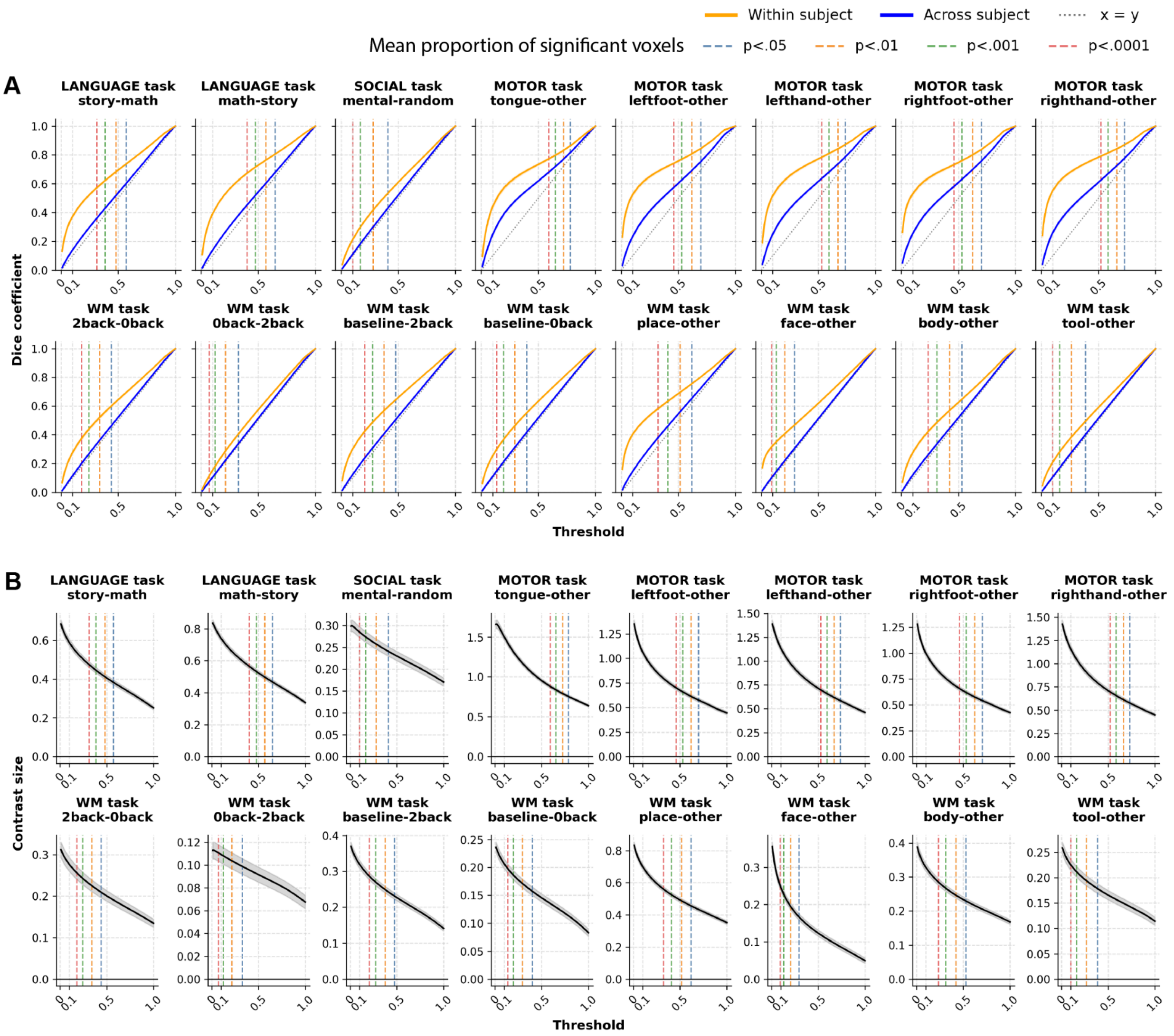


**Supplementary Figure 7.** Robustness of fROI definition across fROI thresholds. (A) Within-subject overlap of fROI definitions as a function of fROI threshold. (B) fROI sensitivity as a function of the fROI threshold.

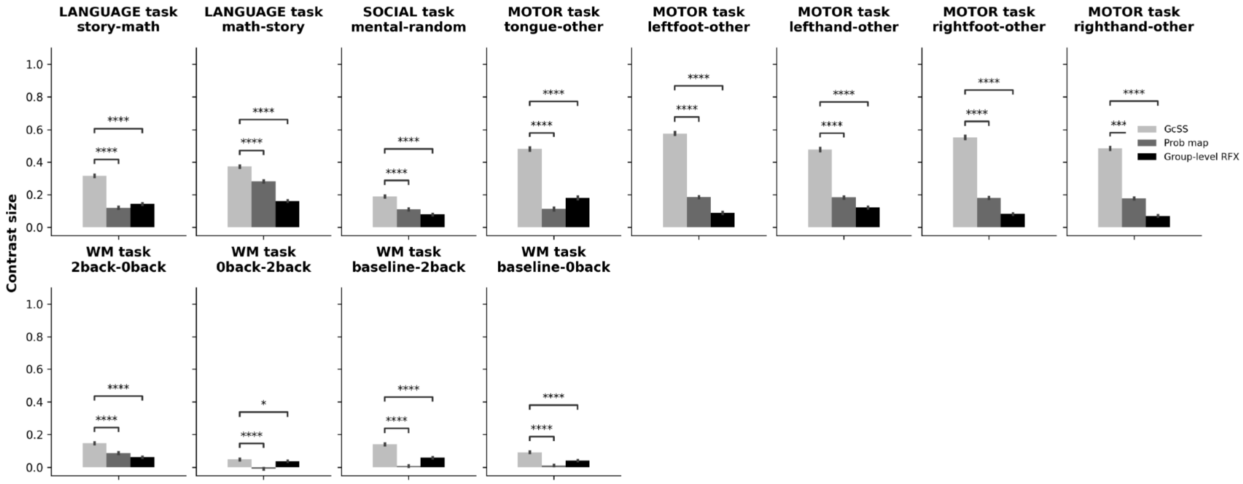


**Supplementary Figure 8.** Effect size for the contrast of interest in ROIs defined using GcSS vs. a probabilistic group-level map vs. the traditional RFX analysis in the cerebellum.

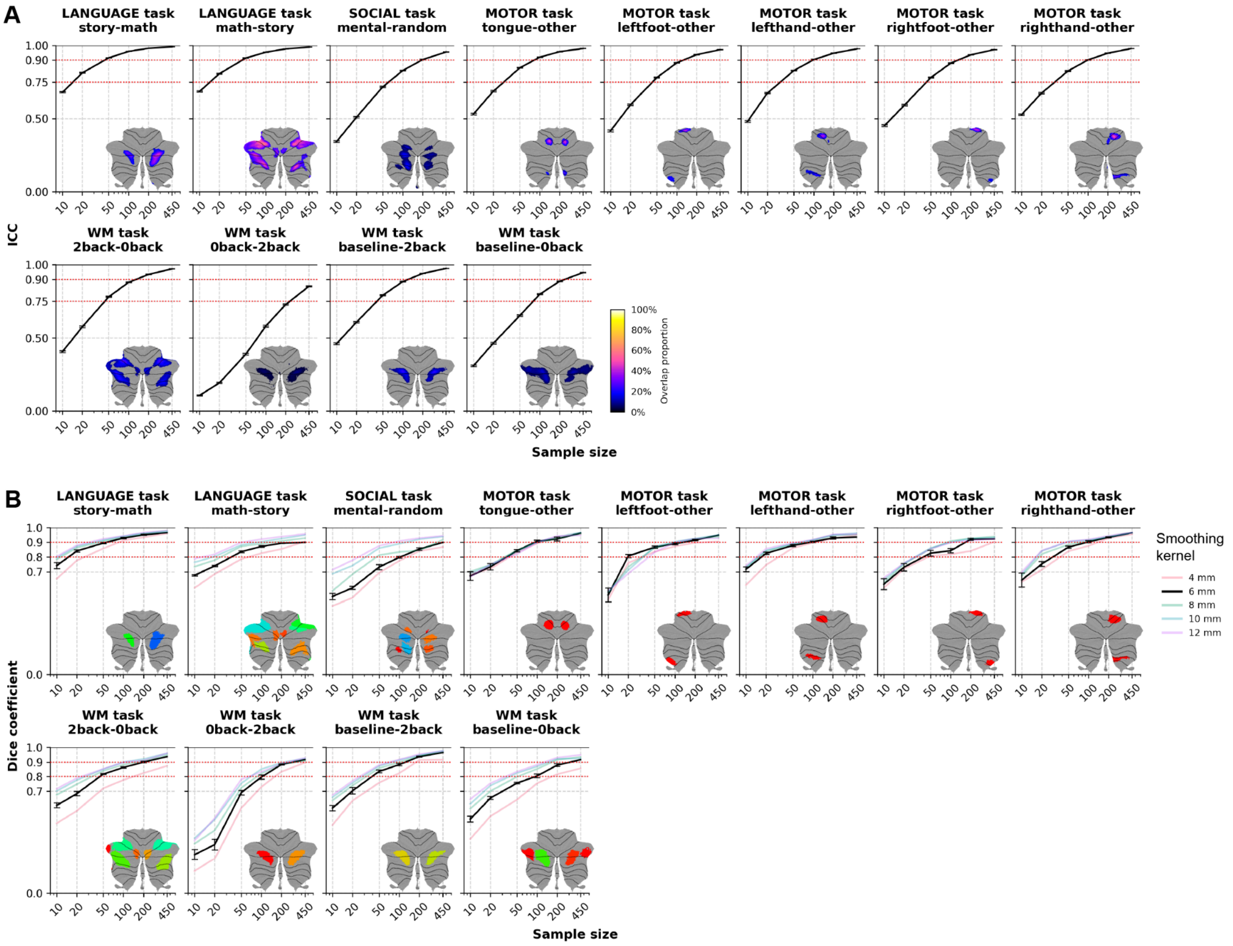


**Supplementary Figure 9.** Reliability of probabilistic maps and group parcels in the cerebellum. (A) Intraclass correlation of probabilistic map estimation over voxels as a function of sample size. Error bars denote 95% confidence intervals. (B) Dice overlap coefficient of group parcels as a function of sample size. Error bars denote the standard error of the mean.

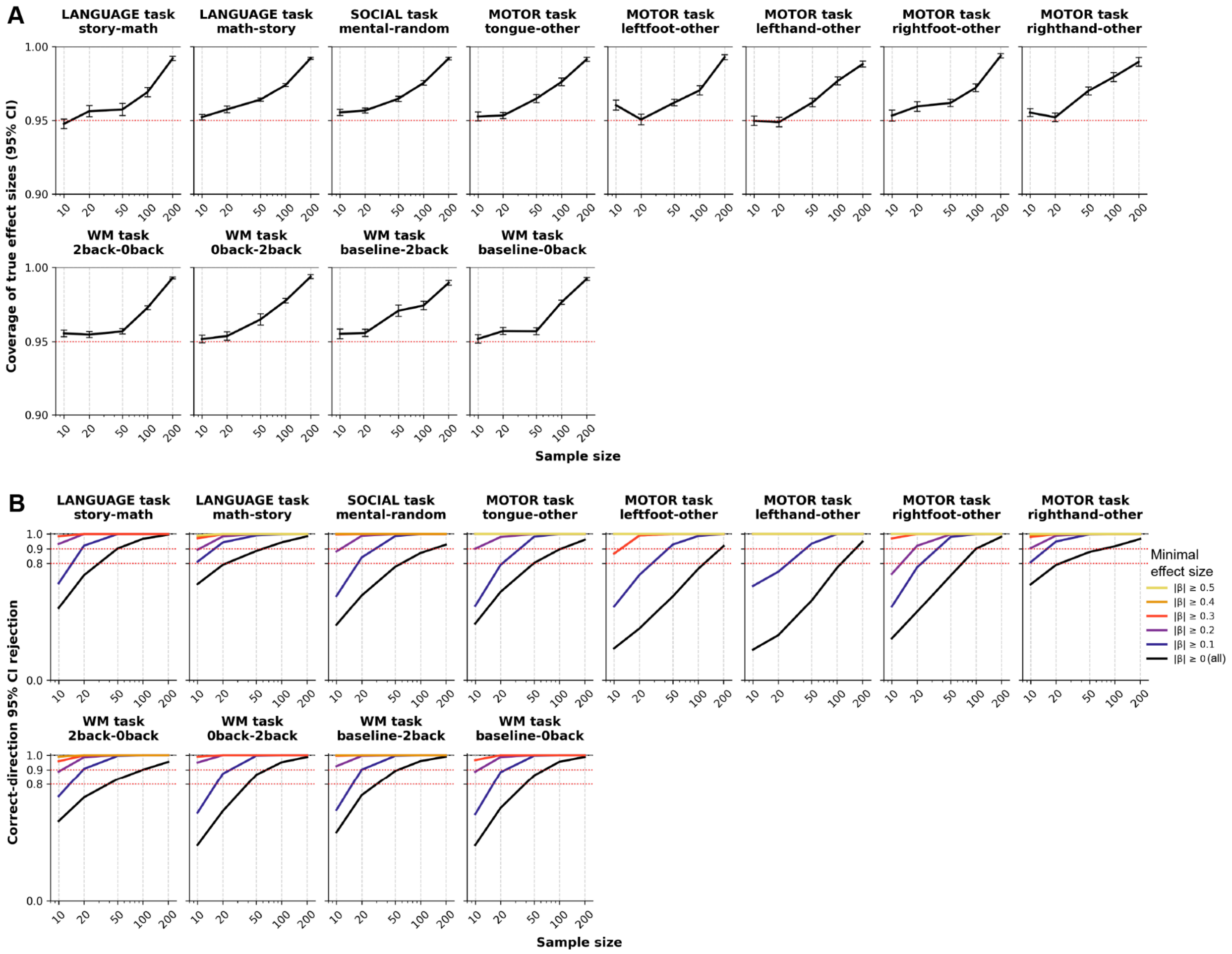


**Supplementary Figure 10.** Validity and power of 95% confidence interval (CI) in the cerebellum. (A) Mean proportion of samples in which the 95% CI includes the ground-truth effect size (computed from all held-out subjects, N > 450), as a function of sample size. Error bars denote standard errors across fROIs and effect conditions. (B) Mean proportion of samples which successfully rejected the null hypothesis for significant effects, as a function of sample size and effect size.

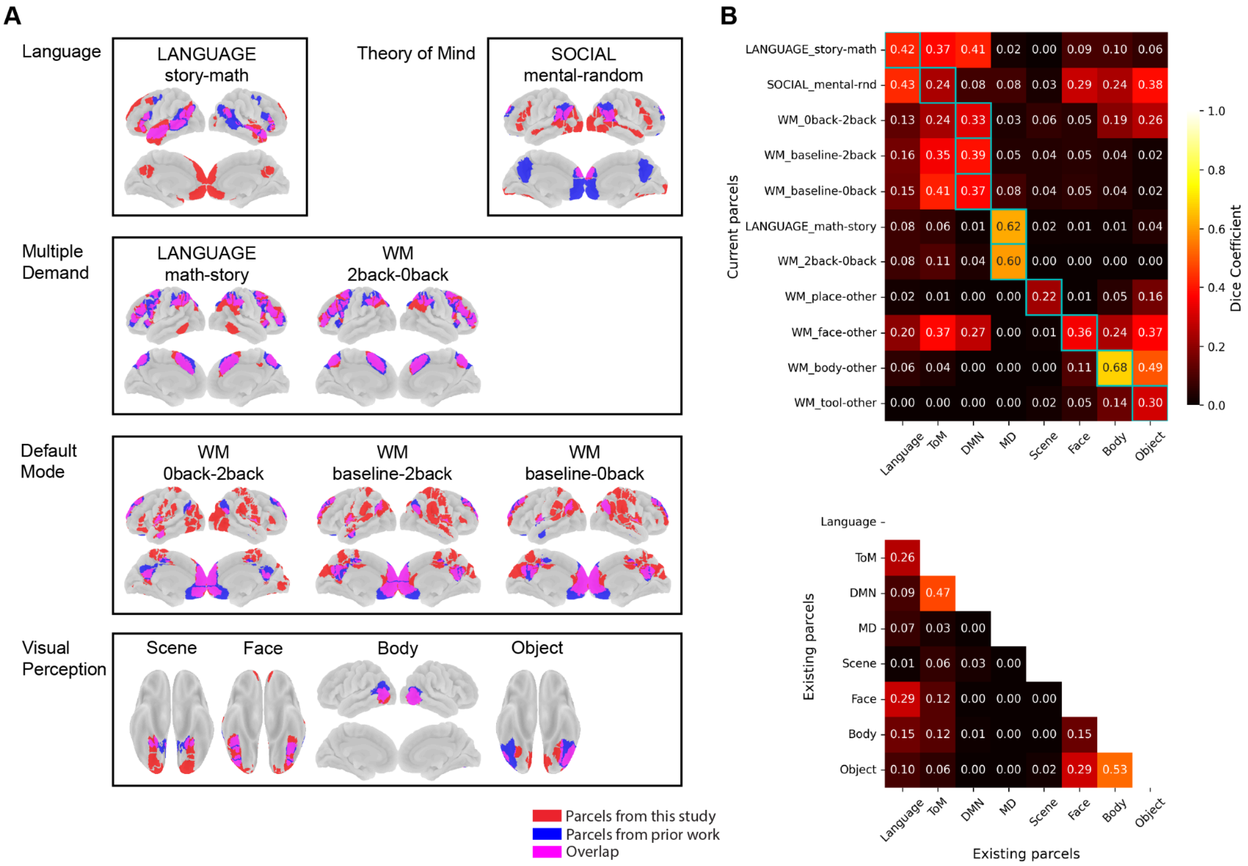


**Supplementary Figure 11.** Cross-dataset comparison of GcSS-derived parcels. (A) Spatial overlap between parcels from the current dataset and previously published GcSS network parcels. (B) Dice overlap between current parcels and reference network parcels. (C) Dice overlap among reference network parcels.

#### Supplementary Note 1

As many fMRI-based analyses are performed after mapping data onto anatomical surfaces, we conducted additional analyses leveraging Human Connectome Project surface data. <TODO: add details later>. The scaling laws of surface-based probabilistic maps are largely similar to those of volume-based maps (**Supplementary Figure 12A**), exhibiting very similar topographic patterns but generally higher overlap values. Surface-based maps show modest increases in probabilistic consistency at smaller sample sizes for most contrasts, most prominently in face perception regions. However, this advantage becomes less pronounced as sample size increases. When the data and group parcels are projected onto the surface, the estimated effect sizes are similar to those estimated from the volumetric data across all fROIs (**Supplementary Figure 12B**).

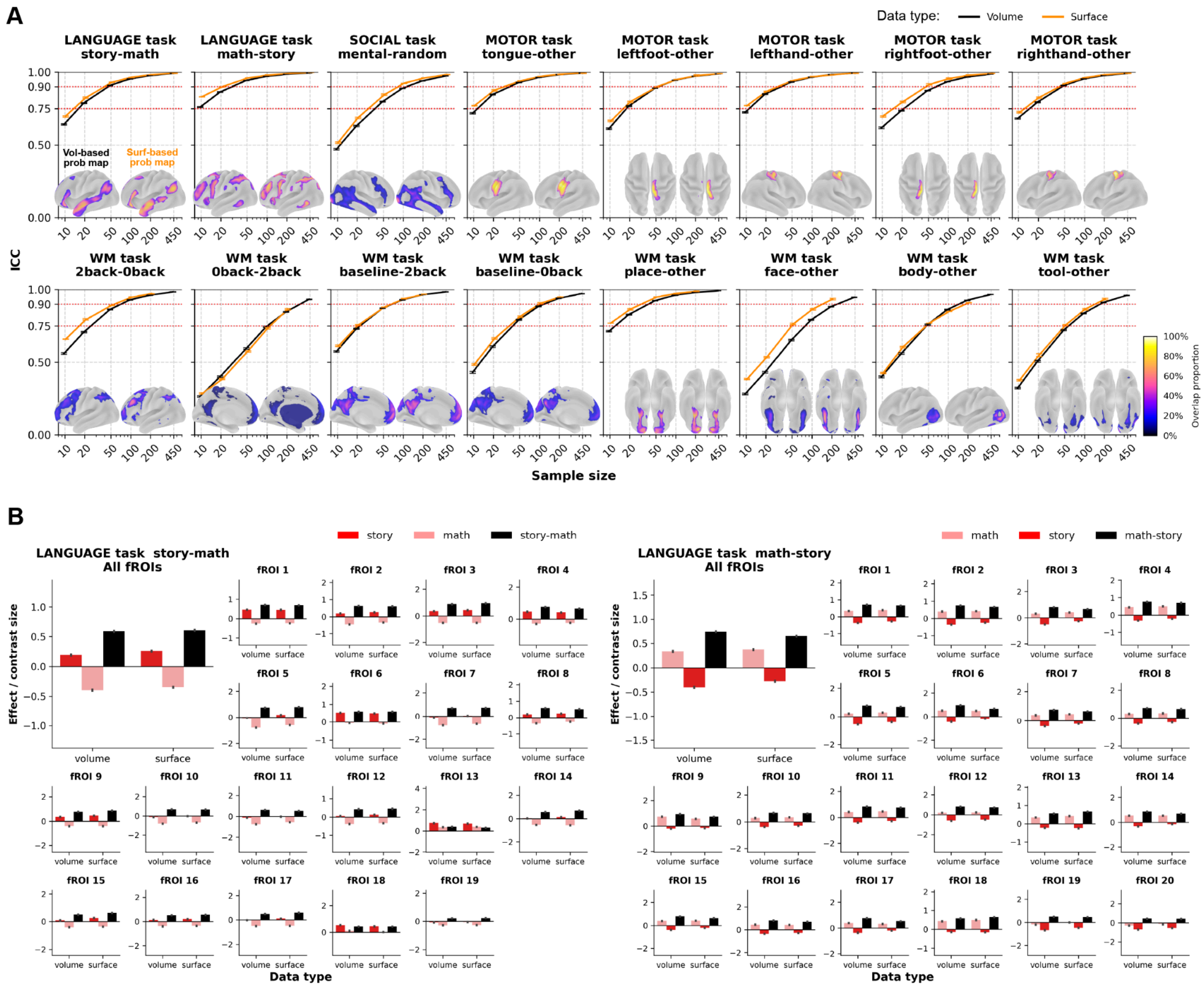


**Supplementary Figure 12.** (A) Intraclass correlation of probabilistic map estimation over voxels as a function of sample size. (B) Example effect sizes estimated on volume data vs. surface data.
